# Horizontal connectivity in V1 : Prediction of coherence in contour and motion integration

**DOI:** 10.1101/2021.09.10.459746

**Authors:** Benoit Le Bec, Xoana G. Troncoso, Christophe Desbois, Yannick Passarelli, Pierre Baudot, Cyril Monier, Marc Pananceau, Yves Frégnac

## Abstract

This study demonstrates the functional importance of the Surround context relayed laterally in V1 by the horizontal connectivity, in controlling the latency and the gain of the cortical response to the feedforward visual drive. We report here four main findings : 1) a centripetal apparent motion sequence results in a shortening of the spiking latency of V1 cells, when the orientation of the local inducer and the global motion axis are both co-aligned with the RF orientation preference; 2) this contextual effects grows with visual flow speed, peaking at 150-250°/s when it matches the propagation speed of horizontal connectivity (0.15-0.25 mm/ms); 3) For this speed range, the axial sensitivity of V1 cells is tilted by 90° to become co-aligned with the orientation preference axis; 4) the strength of modulation by the surround context correlates with the spatiotemporal coherence of the apparent motion flow. Our results suggest an internally-generated binding process, linking local (orientation /position) and global (motion/direction) features as early as V1. This long-range diffusion process constitutes a plausible substrate in V1 of the human psychophysical bias in speed estimation for collinear motion. Since it is demonstrated in the anesthetized cat, this novel form of contextual control of the cortical gain and phase is a built-in property in V1, whose expression does not require behavioral attention and top-down control from higher cortical areas. We propose that horizontal connectivity participates in the propagation of an internal “prediction” wave, shaped by visual experience, which links contour co-alignment and global axial motion at an apparent speed in the range of saccade-like eye-movements.

## Introduction

Low-level visual perception is often described as a “pop-out” holistic process, elevated at the state of awareness, independently of behaviorally directed attention. At the perceptual level, Gestalt rules link stimuli according to feature similarity, spatial proximity or common fate (1, 2). Their neural implementation is thought to occur at various stages of visual cortical processing (3) and best captured, in terms of receptive field (RF) computation, as the combination of smooth “association field” operators in space (4, 5) and in time (6). However, the detailed synaptic implementation and the cortical regionalization of this dual space-time mechanism remain largely unknown (review in (7)). An unsolved issue, which is central to this study, is to determine to which degree “horizontal” connections, intrinsic to the primary visual cortex (V1), are instrumental to the neural implementation of Gestalt laws.

Classically, the consensus is that binding in space and time are processed in separate cortical areas, respectively in V1 for local attributes of form and contour integration, and in MT for global motion (8–10). Although the functional expression of contour-related responses in V1 has been shown to depend on the behavioral attention context and to be task-specific (11), it requires the triggering of a stimulus-driven bottom-up process already detectable in the anesthetized brain (12–14). Local circuits intrinsic to primary visual cortex such as the long-range horizontal connections, are involved in contour integration, but the final binding process is gated and tuned in strength by selective top-down signals conveyed by re-entrant cortico-cortical pathways ((11, 15); review in (16)). In contrast, in terms of hypothetical cortical circuitry contribution, neural correlates of motion binding appear well established in MT (17–19) but far less documented in V1 (but see (20)).

In spite of many intracellular receptive field (RF) studies in cat V1 (21–24), only a few have directly measured the synaptic nature of contextual influence originating from the “silent surround” (25–30). A striking feature from our own observations is that the strength of distal synaptic echoes exponentially decreases, while their latency linearly increases, with the relative eccentricity of the stimulus from the RF center. If intracellular studies quantify synaptic convergence of lateral input, optical imaging, in a converse way, monitors the full network depolarization pattern and dynamics in response to a local stimulus. Intrinsic imaging reveals the spatial extent of the diffusive pattern of intracortical activity, spread across several hypercolumns (up to 6° in cat, in (31)). Mesoscopic dynamic imaging, using voltage sensitive dyes with a temporal activation precision in the millisecond range, report depolarizing wavefronts propagating across the superficial layers of V1, observed both in the ongoing and visually driven states (non human primate (NHP) (32–36); cat (37–40); rat (41)). Their speed range is comparable across mammalian species and matches remarkably closely that inferred from our intracellular recordings in the cat (between 0.05 to 0.50 mm/ms in (26)).

Intracellular recordings provide a unique way to dissect out the diversity of synaptic input connectivity pattern across individual V1 cells. Through selective averaging, they allow to infer the mean functional kernels of feedforward and lateral synaptic inputs composing the V1 receptive field. A prior study in our lab (29) proposed a new feature-specific averaging method, while keeping the mapping stimulus geometry invariant across cells in a receptive-field-centric framework. This was done by realigning each of the individual subthreshold maps relatively to a common origin and “ 0° ” reference axis, corresponding respectively to the center and the preferred orientation/direction of each recorded RF. The averaging of 25 RF maps demonstrated that, irrespective of single cell RF diversity, the mean synaptic connectivity kernel of V1 neurons receives predominantly iso-oriented inputs from neighboring collinear V1 hypercolumns. A remarkable feature is that its spatial organization pattern exactly mirrors the perceptual “Association Field” for collinear contours in human psychophysics (4, 42).

The extensive intracellular study on dynamic Center-Surround interaction we present here - on a larger cell sample (n=65) - is based on the same premise. It seeks to maximize the paired neighbor-to-neighbor interactions between Surround and Center and quantify a novel form of contextual control on the spiking responsiveness of V1 cells. For this purpose, we have designed here a variety of multi-stroke apparent motion (AM) sequences of specific speed and spatial anisotropy, which, under precise timing and local feature co-alignment conditions, allowed us to control the temporal phase between horizontal and feedforward inputs (Figure 1A). Two sets of visual protocols were applied separately in two distinct sets of cells with complementary objectives. The “**cardinal**” protocol (Fig. 1B), was designed to : i) optimize the visibility of synaptic inputs originating laterally from the “Near” RF surround, outside the spiking “minimal discharge field” (MDF), but potentially overlapping with the subthreshold depolarizing receptive field (SRF); ii) probe for a possible axial bias in the preferred recruitment of contextual Surround information integration, using 2-3 stroke AM sequences along the two main (length/width) axes of the RF). The second series of protocols, termed “**radial**” (Fig. 1C), was designed to minimize the risk of spurious feedforward contamination when probing the locations closest to the RF borders (position D1 in Fig. 1A). Accordingly, the synaptic feedforward cortical imprint was no longer limited to the spiking discharge field (MDF), but included as well the surrounding depolarizing SRF evoked by point-like stimuli during sparse noise mapping. Conversely, this new requirement restricted the Surround stimulation to “Far” regions outside the SRF. To compensate for potentially weaker Surround responses, we increased the number of AM strokes and imposed radial symmetry in order to recruit simultaneously more sources in the RF periphery at the same relative Surround eccentricity.

**Fig 1.**
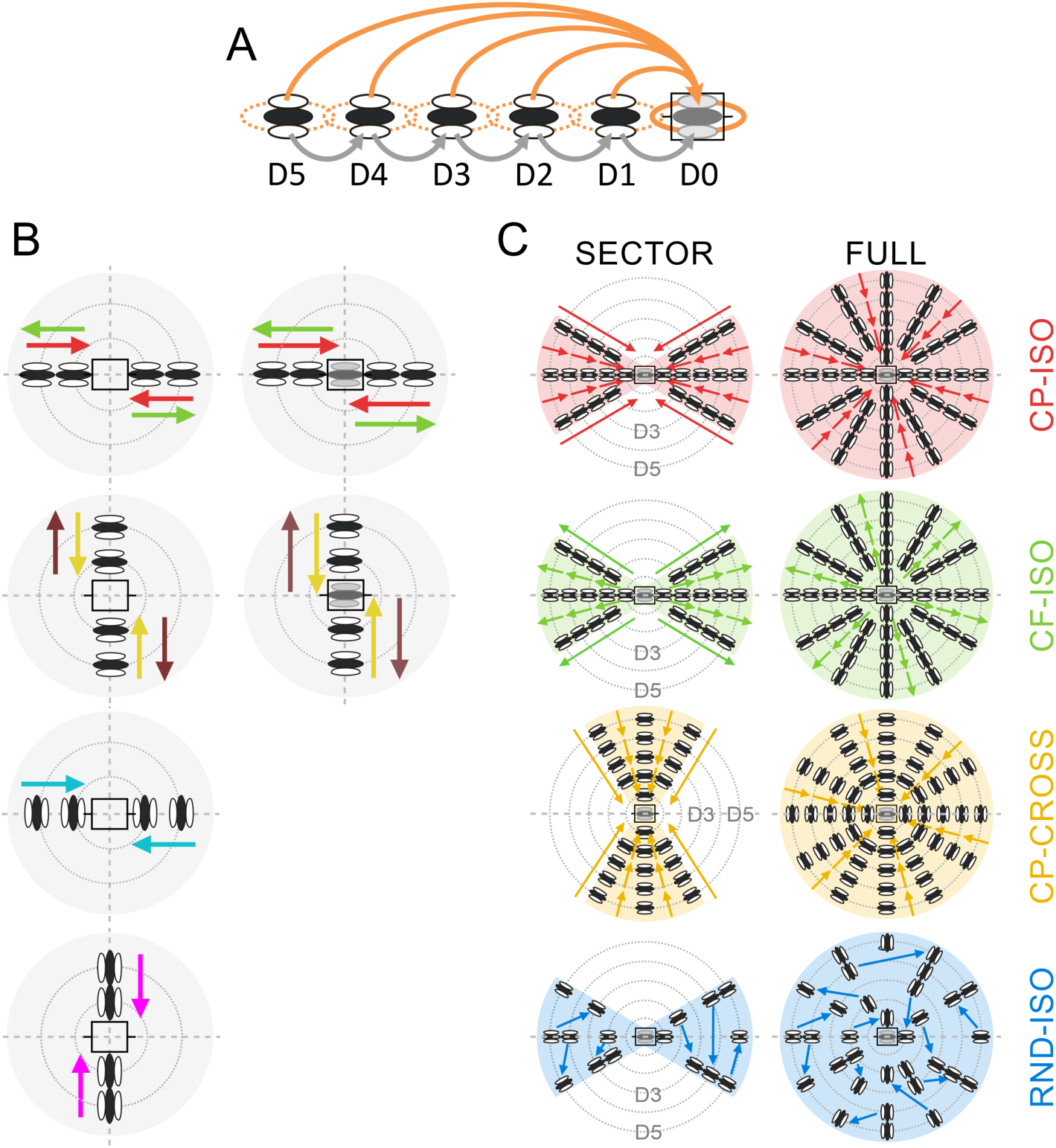
Working Hypothesis and Visual Stimulation Protocols. ***[A] : Working Hypothesis :*** the AM sequence is designed so as to synchronize the synaptic activation time of horizontal volleys, evoked by the sequential presentation of 2 to 5 Gabor patches (GP) in the Surround, with the feedforward activation of the RF Center (shaded rectangle in the D0 position). The Surround GPs, regularly spaced from “Far” (D5) to “Near” (D1) periphery along the apparent motion path (horizontal axis), are flashed in succession at high contrast. The test reference GP, terminating the AM sequence, is flashed at low/medium contrast in the RF center (D0). Each orange arrow represents the lateral propagation of the presynaptic volley elicited by each distal GP. Other synaptic recruitment paths potentially also contribute also to Surround-Center modulation, in particular cascades of proximal neighbor-to-neighbor links (grey arrows). ***[B] “Cardinal” Protocol :*** the two columns illustrate the exploration of the two cardinal axes (horizontal for the RF orientation preference; vertical for the RF width axis) with 2 or 3-stroke AM sequences of iso-oriented stimuli (“collinear”, first row; “parallel”, second row) or cross-oriented stimuli (two bottom rows, same motion paths but the local inducer orientation is now orthogonal to the RF orientation preference). ***[C]: “Radial” Protocol*** : the two columns illustrate the various AM flow patterns (individual rows), either funnelled along a given motion axis (**SECTOR**, left), radially contracting (CP: centripetal) or expanding (CF: centrifugal), or across the full Surround (**FULL**, right). In the SECTOR condition, two opposite motion axes are symmetrically explored, aligned either with the RF orientation preference (horizontal axis (± 30°) in rows 1, 2 and 4) or with its width axis (vertical axis (± 30°), row 3). In the FULL condition, all directional motion axes (discretized by 30° steps) are stimulated concurrently and intersect the RF center (horizontal grey icon). From top to bottom, each row illustrates one specific configuration of AM flow: 1) centripetal collinear flow (**CP-ISO**) from Surround to Center (red, row 1); 2) centrifugal collinear flow (**CF-ISO**) from Center to Surround (green, row 2), 3) centripetal cross-oriented flow (**CP-CROSS**) across the RF width axis (gold, row 3); 4) randomized collinear pattern (**RND-ISO**) where each successive GP location has been randomized in space and time (blue, bottom row). Note that in all the conditions other than CF-ISO, the last GP of the sequence is flashed with the optimal orientation in the RF center (D0). See <u>Text</u> for details.

Our working hypothesis posits that the spatio-temporal optimization of the control of cortical responsiveness should be reached <u>where, *in space*</u>, the local orientation of each Surround GP matches that of the recorded RF (as expected from the “Association Field” profile), and <u>when, *in time*</u>, the phase of the stimulation of each surround node compensates for the lag of the postsynaptic response, due to the dependency of the PSP latency on the eccentricity relative to the RF center (Fig. 1A, “latency basin” in Fig. 2A). Our functional prediction is that a latency shortening and an overall cortical gain amplification of the composite evoked PSP should be observed when the global flow speed of the AM sequence synchronizes the respective synaptic impacts of the lateral and feedforward drives onto the intracellularly recorded cell.

**Fig 2.**
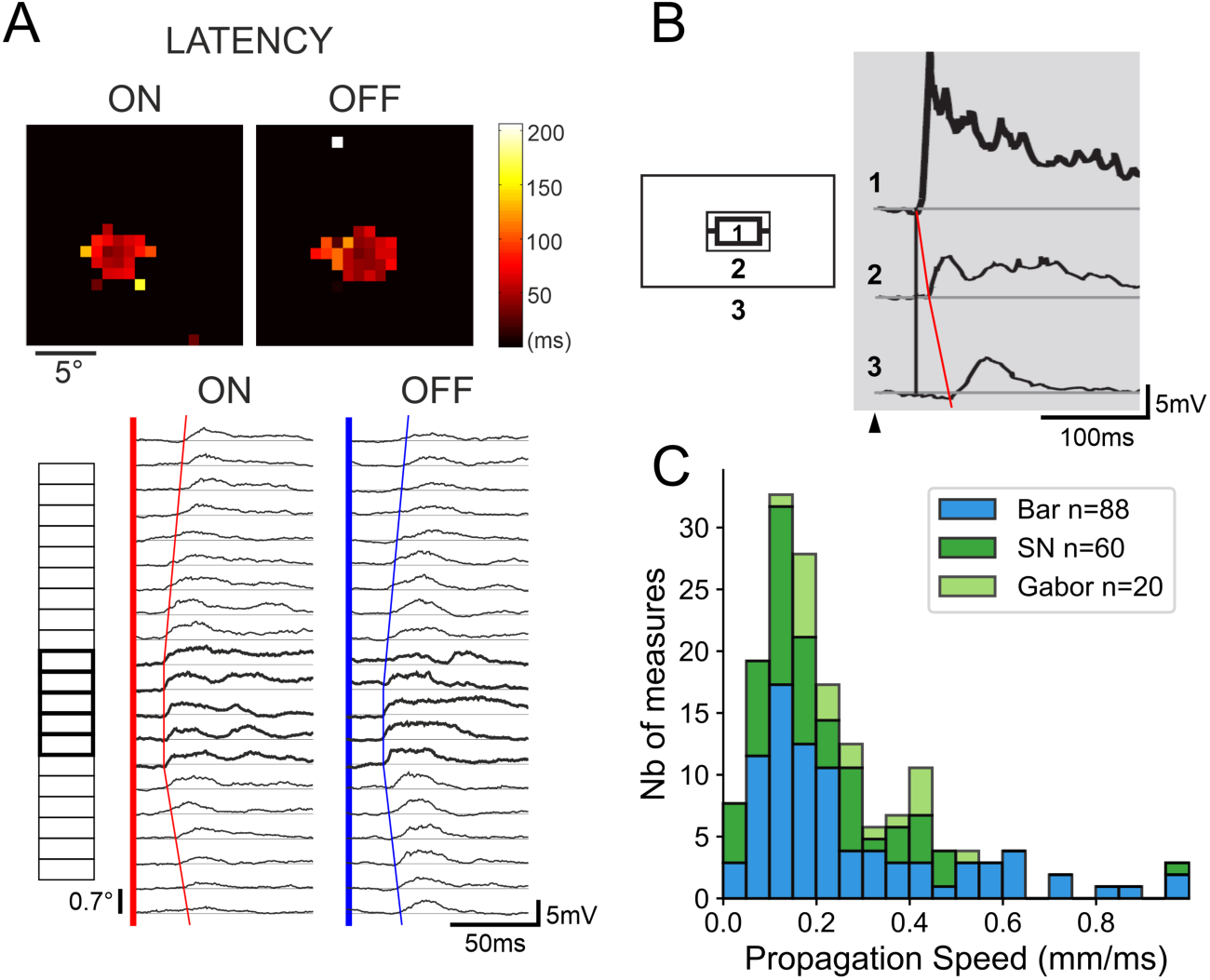
A synaptic view of visual V1 receptive fields. ***[A] : Latency basin of synaptic responses relative to the RF Center :*** Top left, spatial maps of the ON- and OFF- Depolarizing Fields latencies using sparse noise (SN) mapping. Heat scale: color code for latency values (from 0 (black) to 200 ms (yellow)). Bottom left, 1D-mapping of ON- and OFF- PSP responses in the same cell, evoked by long bars (7.1° x 0.7°) flashed at the optimal orientation and positioned at different eccentricities (ordinates, ranging from -4° (bottom) to +6° (top)) across the RF width axis. Thick boxes (along the left ordinate axis) designate the positions which are in overlap with the SRF mapping using SN (above, top left). Red (ON) and blue (OFF) linear regressions illustrate the 1D-latency basin profiles of Surround-Only subthreshold responses. The linear fits of latency versus eccentricity give apparent horizontal speeds of propagation estimates of 0.18-0.38 mm/ms for ON- (red, left) and OFF- (blue, right) responses. ***[B] Dependency on spatial summation*** (adapted from (26)) : From top to bottom, phase-reversal (2 Hz) responses to disk [1] and annular gratings [2 & 3], for increasing inner border eccentricities from the RF center (0° in [1], 5.6° in [2] and 10.3° in [3]). Note the decay in response strength and the increase of onset latency in Surround-Only responses [2 & 3] with eccentricity (reaching 32 ms for the “Far” Surround annulus). ***[C] Apparent Speed of Horizontal Propagation (ASHP) distribution*** : Stacked data histograms from (26) using long bars (1D-mapping, blue) or sparse noise (green), and from this study (Gabor, light green). Note that, independently of the probe stimulus, most AHSP values inferred from Surround latency basin slopes range between 0.05 and 0.60 mm/ms.

This study addresses the importance of the Surround context dynamics relayed laterally in V1 by the horizontal connectivity, in controlling the timing and the responsiveness gain of the cortical response. It also gives evidence for neural correlates of <u>“filling-in”</u> responses at the earliest stage of visual cortical processing, i.e. the primary visual cortex. Remarkably, these responses are shown to be specific of co-alignment, and their timing anticipates the occurrence of the same orientation feature in the RF center, even in the absence of feedforward activation. We infer, from a detailed contextual synaptic interaction analysis, a conceptual framework which accounts for the neural correlates in V1 of specific Gestalt-like visual configurations, linking local (orientation) and global (contour integration, motion) features.

## Material and Methods

### 1. Animal preparation and electrophysiological recordings

All experiments were performed in anesthetized (alfaxalone, Alfaxan®, 2.5 mg/kg/hr) and paralyzed (pancuronium Pavulon® (0.1 ml/kg/hr) or rocuronium Esmeron®, 4 mg/kg/hr) adult cats of either sex, in accordance with the American Physiological Society’s Guiding Principles for the Care and Use of animals. This study was performed in institutionally approved animal breeding and experimental facilities (CNRS Gif Campus : Central Animal Care facilities ; authorization # D91-272-105). The scientific project was approved by the French Ministry of Higher Education and Research (MESR) (# 3414-201603111-3088501) and monitored by the local governmentally approved CNREEA Ethics Committee (CEEA – 059 : *Comité d’éthique en matière d’expérimentation animale*, Region Paris Centre et Sud). Experiments where continuously supervised by a veterinary DVM. Every effort was made to minimize stress and suffering. Anesthesia depth, analgesia level and the general physiological state stability were continuously monitored from the recording and analysis of EKG and extradural ECoG signals. End-tidal CO2 was maintained between 3.5 and 4.5%, and body temperature was regulated around 38°C throughout the experiment. Once the animal’s head was fixed to the stereotaxic frame, and the craniotomy performed, we checked again the presence of a slit pupillary state, indicative of adequate anesthesia during the surgery phase. Then, atropine (1%) and phenylephrine (5%) were applied to both eyes to dilate pupils, block the accommodation and withdraw nictitating membranes. Artificial pupils and corrective lenses were used to protect the eyes and focus the backprojection of the eye fundus on the stimulation screen. At the end of the recording session (48 to 80 hours), the animal were sacrificed using either an overdosage of pentobarbital (180 mg/kg, iv) or an intra-cardiac perfusion performed after alfaxalone injection rate was increased high enough to induce a flat EEG.

*In vivo* intracellular recordings of cortical cells were performed in the primary visual cortex (area 1. 17) using sharp Glass electrodes pulled from 1.5-mm-thick borosilicate capillaries (World Precision Instruments Inc.). Electrodes were filled with a 2M potassium methyl sulfate (pH ranging from 7.2 to 7.4), 4 mM KCl solution for compensating the tip potential, and 1.5% biocytin. Electrode resistances before impalement ranged between 70 and 130 MΩ. Cells were recorded in current clamp bridge mode using an Axoclamp 2A (Axon Instrument inc.) or a SEC-05X (NPI inc.). The average resting potential was between -62 to -78 mV (mean ± σ = -66,8 ± 8,3 mV) with no or a slight holding current (<= -0.5 nA) when needed. Signals were filtered and amplified with a CyberAmp380 (Axon Instrument inc.) and digitized at 10 kHz by an ITC-1600 (InstruTECH®) card. Based on recording depth and the laminar location and morphological identification of fourteen biocytin labeled cells, the overall sampling was distributed from supra- to infra-granular layers. However, in view of the limited number of labelled cells (experimentally constrained by gliolysis increase with the length of recording) and a slight bias for recordings in deep layers, no laminar study was done in this reduced sample.

### 2. Visual stimulation

For the “cardinal” protocols, visual stimuli were generated, using the ELPHY in-house software (G. Sadoc, UNIC-CNRS), on a CRT P31 green monochrome monitor positioned at 57 cm (1 cm = 1° of visual angle) at refresh rate of 60 Hz (19 inches, 640 x 480 pixels). For the “radial” protocols, we used gamma-corrected LCD green monitor (27 inches screen, 1920 x 1080 pixels) at 28.5 cm from the cat’s retinal plane (0.5 cm = 1° of visual angle) at a refresh rate of 144 Hz .

#### 2.1. Initial quantification of RF optimal features

The receptive field (RF) position of each recorded neuron and its ocular dominance were initially established using moving light bars. Stimulation was done through the dominant eye for the rest of the experiment. Subthreshold ON and OFF (SRF) sub-regions and spike minimal discharge fields (MDF) were then mapped by forward correlation of visual responses to a random “sparse noise” (SN) protocol : white (40 cd/m^2^) and dark (1 cd/m^2^) non-overlapping squares were presented one at a time for 16-34 ms against a grey background (20 cd/m^2^) in each position of the exploration grid (8×8 or 20×20 pixels) centered over the coarse RF region. In some cells, an additional 1D mapping was done with an optimally oriented thin long bar, flashed randomly at various eccentricities across the RF width (Fig. 2A).

Once the RF location defined, the preferred orientation, phase and spatial frequency of the cell were determined using a dense Gabor Noise (GN) stimulation protocol (43) : flashed (16-34 ms) Gabor patches (GPs, adjusted in size to the s-DF extent) were randomly presented at six different orientations (with 30° steps), four phases (0°, 90°, 180° and 270°) and five spatial periods (ranging from 0.1 to 4.9 cycles per degree of visual angle). For each cell, the parameters of the Gabor patch eliciting the strongest response in the RF center were retained to compose a template of preferred features of the inducer (independently of its position) in the subsequent apparent motion (AM) sequences. In order to evoke a sizeable synaptic “horizontal” response, the GP flashed in the Surround were always presented at high contrast (mean ± σ = 92.5 ± 11.5 %; range 75-100%), whereas the feedforward central stimulus was flashed in the MDF either at low or medium contrast (mean ± σ = 40,6 ± 24.5 %; range (25-100%)).

#### 2.2. “Cardinal” Apparent Motion (AM) Protocol

In order to assess the dependency of the center-surround interaction on local orientation and global motion, we first designed two/three-stroke center-surround AM protocols for which the Gabor patches were flashed along the two “cardinal” RF axes defined by the RF preferred orientation and width (Fig. 1B). We looked at the modulatory effect that a two-stroke activation of the “silent” surround would produce (along each of these axis) on the response to a test stimulus flashed within the MDF (the third stroke). Randomly interleaved three-stroke stimulation sequences were applied moving away from the receptive field center (“centrifugal” (CF) condition, green arrows in Fig. 1B), or from the surround to the center (“centripetal” (CP) condition, red arrows) from one side of the RF at a time.

The size of the Gabor patch mask was set at 150% of the length of the spiking discharge field while the standard deviation of its Gaussian envelope was set at 20%. The stimulus inter-node distance was of 120% of its length, often leading the more proximal GP to partially encroach on the border of the impulse SRF. The orientation of the local inducer could be defined two ways, either in relation with the RF orientation preference, or in relation with the AM motion axis. These two analysis contexts will be considered separately in the Results. Here, by sake of uniformity with the “radial” protocol, the ISO- and CROSS-conditions are defined relative to the chosen AM axis. The complete set of Surround-Only stimulations (Fig. 1B, left column) included 8 conditions [CP- ISO_main-axis (red arrows), CP-ISO_width-axis (magenta), CP-CROSS_main-axis (cyan), CP- CROSS_width-axis (gold), CF-ISO_main-axis (green), CF-ISO_width-axis, CF-CROSS_main axis, CF- CROSS_width axis (brown)] X 2 direction conditions (one side at a time)], to which 2 control conditions (1 Center-Only + 1 Blank trial, where no stimulus was presented) were added. Hence a total of 18 conditions was completed in 13 cells. Center-Surround interactions were tested in 23 cells with full AM sequences terminating in the RF center (Fig. 1B, right column), under 10 conditions [4 conditions (CP-ISO_main-axis, CP-CROSS_width-axis, CF-ISO_main-axis, CF-CROSS_width-axis) X 2 directions (one side at a time) + Center-Only stimulation + 1 blank]. For each study (Surround-Only; Surround-Center), the different stimulus conditions were interleaved randomly in each trial block. Recordings lasted for 20 blocks.

Given the constraint of the monitor distance (57 cms), its refreshing rate (60 Hz), and the fixed duration of the static Gabor inducer presentation (16 ms), individual cell AM speed values only depended on the size of the recorded RF which was used to define the sequential interstimulus step distance. Accordingly, AM speeds were preset in the suitable range of values to compensate for the dependency of visual PSP latency with relative eccentricity (26, 29).

#### 2.3. “Radial” Apparent Motion (AM) Protocol

This series of protocols was done with a larger (27 inches) and faster screen (144 Hz), using a shortened viewing distance (28.5 cms), which allowed a more extensive exploration of the “Far” surround. The visual field was paved with a grid composed of 5 concentric rings of increasing eccentricity centered on the RF covering up to 25° in the surround (Fig. 1C). To the difference of the “cardinal” protocol, where stimulus size and spacing were defined relatively to the spiking MDF size, the distance between each outer ring and the patch size were adjusted in this new protocol to the full extent of the subthreshold depolarizing receptive field (SRF) mapped with impulse stimuli. This change in the RF core definition, from that used in the “cardinal” protocol condition, was done to prevent, on a cell-by-cell basis, a direct feedforward contamination by the stimulation of the most proximal Surround location (position D1 in Fig. 1A).

Additionally, in the “radial” protocol, we designed Apparent Motion sequences whose spatio-temporal coherence was parametrized to dissect the dependency of the inducer orientation relative to the motion path, as well as the spatio-temporal congruency within the AM flow (see Results). For each recorded cell, the AM sequence of interest was the centripetal iso-oriented configuration (**CP-ISO**, top “red” row in Fig. 1C), where the local inducer orientation was either co-aligned along the preferred orientation axis of the recorded RF (SECTOR configuration, left column in panel 1C) or along the global motion axes (FULL configuration, right column). Three supplementary visual AM configurations were tested, in which the spatio-temporal structure of the local Gabor patch (GP, local inducer) sequence was altered, while keeping the overall stimulus energy distribution unchanged: i) in the <u>first control</u> condition, namely the centrifugal iso-oriented AM sequence (**CF-ISO**, second “green” row from the top, Fig. 1C), the temporal order of the sequential presentation of the co-aligned GPs was reversed, resulting in a centrifugal (CF) flow from Center to “Far” Surround locations; ii) the <u>second control</u> was designed to study the contextual impact of the local GP orientation relative to the motion axis. In the centripetal cross-oriented AM condition, the local orientation of the inducer was orthogonal to the motion axis (**CP-CROSS**, third “yellow” row from the top, Fig. 1C); iii) in the <u>third control</u> condition, the same retinal space (SECTOR or FULL) as in the CP-ISO condition of interest was stimulated with the same GPs (i.e., with the same number, features, and energy distribution at each step of the AM) but the coherence of the flows was randomized both in space and time (**RND-ISO**, bottom “blue” row). For each block, each stimulus location was visited only once, and the last stimulus location was the RF center. Apart from the RND-ISO condition (Fig. 1C, 4^th^ row), the repeated sequences were identical across blocks (pseudo-randomization).

The stimulus set in the “radial” protocol - where all contextual conditions were explored - was composed of 16 dynamic apparent motion sequences (4 conditions (CP-ISO (red arrows), CF-ISO (green), CP-CROSS (gold), RND-ISO (blue)) X 2 trajectories (with or without Center stimulation) X 2 configurations (SECTOR and FULL), and 21 stimulation types where Gabor Patches were flashed in isolation (5 peripheral locations X 2 conditions X 2 configurations + 1 GP flashed in the Center). In each block of trials, each stimulus was presented only once. Additionally, 6 “blank” trials were also included in each block. The resulting set of 43 stimulations was presented in random order in each block. Recordings lasted for 19-50 blocks (median 50).

In order to optimize synaptic summation between Surround and Center stimulation, the AM sequence speed was function of the spatial extent of the SRF and of the stroke duration of Gabor patches and adjusted cell-by-cell in order to fit the latency basin of “Far” surround response latencies. In the “radial” protocol where AM speed and stroke duration could be independently parametrized, the stroke duration was in the 30 ms range (± 9 ms). It was 16.6 ms in the “cardinal” protocol.

### 3. Intracellular Data analysis

#### 3.1. Data formatting

Spikes were threshold-detected in the raw traces and replaced by alpha functions in the membrane potential (Vm) traces (α(t)=a*t*exp(-t/τ)), where “a” is the slope at the peak of the second derivative of the rising phase, and τ is the half-width of the spike), as done in previous studies (29). Resulting traces were band-pass filtered (0.1-300 Hz) and peristimulus triggered waveform (PSTW) were averaged across trials and conditions. Prior to any statistical analysis, all traces were down-sampled to 1 kHz and smoothed using a sliding average window of 7 ms. For spikes, individual peristimulus time histograms were smoothed using a Gaussian window (σ= 3 ms). For a given trial block containing all conditions, the corresponding mean activity (average Vm and Spiking discharge rate) was subtracted from the raw PSTWs and PSTHs.

#### 3.2. Response quantification and statistical significance

For the “cardinal” protocol, the control Z-statistics for Vm (mean and standard deviation) were computed with a 4 kHz sampling rate over a 100-ms sample of ongoing activity preceding the stimulation. The response waveforms were thresholded in amplitude above the 95% one-sided confidence interval. The “integral response” was defined as the integral value of the thresholded depolarizing waveforms over a predefined response integration window (0-250 ms). For each cell, Wilcoxon paired signed rank (WPSR) tests were applied to compare cell-specific responses (n=23) evoked for two distinct contextual conditions. The contextual modulation was quantified by a facilitation/depression index given by the ratio between the significant integral responses observed in the center-surround vs center-only conditions (or between two different contextual conditions of interest). Similar processing was applied at the spike rate level. Twenty-three cells constitute the population used for the Results section for the “cardinal” protocol.

For the “radial” protocol, since a larger spectrum of stimulus conditions was used, more extensive statistical analyses were done on the individual cell’s responses, without or with selection criteria. All significance tests fell into two categories: quantifying the significance of a response to a stimulus or comparing data statistics between two conditions. To do the former, we relied on the activity during blank trials, which provides the null distribution for visual responses (i.e., the activity that would occur should the cell not respond to the stimulus, which was our “null” hypothesis). Consequently, for each cell, the average ongoing synaptic or spiking activity recorded during “blank” trials was subtracted point by point from each evoked PSTW/PSTH trial activity. This was done to remove, on a cell-by-cell basis, any experimental artifact or activity change common to the blank and stimulations, while avoiding a loss of information on the Vm distribution statistics. This preprocessing explains why the confidence intervals obtained from the permutation test are centered on 0 when evaluating significance regarding spontaneous activity (and why there are negative spikes). For studying the dependency on the stimulus context, we used cell-by-cell paired comparison between distinct tested conditions. Statistical analysis was then done using a randomization permutation test with 10^4^ repetitions.

From these computations, we extracted the null hypothesis waveform envelope for a given confidence interval at each point in time. Only cells with a response to the Center-Only condition significantly larger than the blank (p < 0.01) in the interval 0-to-120 ms (“0” being the onset of the last stimulation in the center when the RF was actually stimulated, “120”ms integrating most of the feedforward onset and offset responses) were considered for further analysis. For PSTW and PSTHs, respectively thirty-seven and twenty-two cells were retained after this initial screening and constitute the population used for the Results section for the “radial” protocol. In the remaining fifteen “nonspiking” cells, analysis was restricted to subthreshold activity.

#### 3.3. Quantification of the contextual change in response gain and phase

The concept of “gain control”, dominant in cortical physiology or psychophysics, tends to reduce information transfer functions to simple scalar static ”gain” measurements. It consequently ignores the kinetic aspects of cortical transmission modulation. The intracellular recordings approach developed here allows to compare both the gain and the phase (or advance/delay as analyzed here) of the impulse response (e.g. the transfer function in linear system theory) in various contextual conditions. When using this engineering terminology, we make the simplifying assumption that the test stimulus presentation is short enough in duration and localized enough in space, so that the feedforward test stimulus response gives an approximation of an impulse response in space and time.

We quantified the time-course and the strength of the contextual control of the cortical transfer function by measuring the changes in i) subthreshold latency (i.e. the “phase”) and ii) the integral value of the depolarizing postsynaptic potential (PSP) response to the complete AM sequence terminating in the recorded cell’s receptive field (i.e. the “gain”). For the “cardinal” protocol, the latency measure was defined by the onset of the mean PSP reaching significance (Z-test, p<0.05). For the “radial” protocol, we chose to use a more conservative measure to avoid spurious detections of fluctuations in the ongoing synaptic noise. Accordingly, the latency change (Δ latency) was measured at half-height of the Center-Only response amplitude peak, separately for Vm and spikes. For the response gain (Δ response), we measured the change in the PSP depolarizing envelope integral (and, eventually, spiking integral), relative to the Center-Only response, over the whole time-course of the test-response. Here, the rationale was to integrate differences in evoked responses for all possible time relationships between feedforward and lateral input. The same statistical analysis (randomization test, 10^4^ repetitions; p<0.05) was repeated for PSTWs and PSTHs for both criteria. The paired-analysis was replicated across conditions to measure statistical significance between the main condition of interest (CP-ISO) and the other contextual conditions.

### 3.4. Surround-Only responses

For the “cardinal” protocol, statistical significance was assessed using Z-test statistics on response integrals (p<0.05). For the “radial” protocol, we used a permutation test (10^4^ repetitions; p< 0.01). Due to the noisy nature and the weak amplitude of purely lateral responses, we only retained Vm waveforms for which the statistical significance threshold was trespassed for at least 15 consecutive milliseconds in the interval 0-120 ms.

### 3.5. Linear predictions and Dynamic non-linearities

For the “radial” protocol, we determined in each individual case whether the actual Surround-Only responses to dynamic sequences were larger than (or equal to) the sum of the temporally shifted responses to each GP flashed in isolation at each location (reproducing the spatial and temporal ordering of a virtual AM sequence). This linear predictor was in a first step compared to the actual AM dynamic sequences before they reached the RF Center. A non-linearity was detected in cells where the actual build-up of anticipatory activity during AM sequences deviated significantly from the linear predictor. Because GPs, when flashed in isolation in the close vicinity of the SDF, already induced significant depolarizing responses (29), we considered Surround-Only synaptic waveforms as significantly larger than their “static” linear predictor, if they crossed the upper limit of the confidence interval for a minimum duration of 7 consecutive ms in the interval 0-120 ms (permutation test, 10^4^ repetitions, p < 0.05). A second predictor was calculated by subtracting the Center-Only response from the complete AM “Surround-then-Center” response. Note that the Surround-Only component potentially included the synergetic effect arising from the interaction between the successively recruited distal input sources in the Surround. This two-term (Center/Surround) linear predictor was compared to the observed “Surround-Only” AM response to test for non-linear interactions between the integration of the AM contextual information in the “Near” Surround and the flashed feedforward drive.

#### 3.6. Population analysis

In order to compare responses between cells, each individual flow pattern’s specific geometry – defined on a cell-by-cell basis according to the preferred orientation and direction of each cell – was realigned on a common “0°” reference axis (visualized by the semi-horizontal axis pointing to the right in all Figures), regardless of the absolute orientation preference of each cell. No normalization of responses was used in the “cardinal” protocol. In the “radial” protocol, the amplitude of the PSTWs and PSTHs of each cell were first normalized to the peak of the Center-Only response. Before averaging, PSTWs and PSTHs of all the cells were also realigned with respect to the onset latency of their individual Center-Only response. This common origin was defined as the time abscissa of the first point (blue dot) of the average Center-Only response departing from the “blank” response mean by more than 3σ . This process, replicated independently for subthreshold (Vm) and spiking responses, allowed a realignment of all normalized cell-by cell profiles and the calculus of the mean population response profile. Note that the peak value of the average population PSP can be less than the norm (1.0) when the onset-to-peak rise time varies across cells.

## Results

This intracellular study (65 cells) of Center-Surround interactions in V1 of the anesthetized cat is based on two distinct visual protocols, applied in two different sets of cells (see <u>Methods</u>). The first protocol, termed “**cardinal**” (Fig. 1B), recruiting both “Near” and “Far” RF surround, was applied to a first batch of 23 cells. The second series of protocols, termed “**radial**” (Fig. 1C), limiting the Surround stimulation to “Far” regions outside the SRF only, was performed on a second batch of 42 cells. Accordingly, response averages were done separately for the two visual stimulation protocols.

### 1. Probing synaptic responses from the “silent” Surround

#### 1.1. Spatial summation

The “silent Surround” is classically defined as the region outside the minimal discharge field (MDF), where impulse stimuli do not evoke a significant spiking response. However, the intracellular mapping of synaptic responses with sparse or dense noise (SN or DN), devoid of spatial summation, shows a broader subthreshold (non-spiking) receptive field (SRF) extending beyond the border of the MDF, and often invading the opponent contrast discharge field, even in Simple cells (43, 44). This defines a Near-Surround domain, connex to the MDF, where the contribution of purely lateral synaptic input cannot be unambiguously disentangled from subthreshold feedforward activation (25).

The mapping of the minimal discharge field (MDF) with small light or dark squares (0.2-0.9°) showed on average a spiking core field of 1-3° of visual angle. Overall, the size of the impulse subthreshold depolarizing receptive field (SRF) of the cortical neurons recorded in this study ranged from 2.5° to 7.5° (5,1 ± 1,6°), for eccentricities from the area centralis between 1.1° and 8.3° (4.0 ± 1.6°). This initial mapping was used to define, in a conservative way in each cell, the retinotopic imprint of the feedforward input, whether it produced a spike (MDF) or only a depolarizing subthreshold response (SRF).

In contrast, elongated light bars (5-12° length) flashed across the RF width axis, which increases input spatial summation, elicited synaptic responses, in the same cell, originating much further away from the RF center (7.1° for bars against 2.5° for SN in the example cell in Fig. 2A). Using annular gratings at the optimal phase and orientation, with still larger spatial summation, our lab previously demonstrated distal synaptic responses originating beyond 10° of relative eccentricity (Fig. 2B, taken from (26)). Pooling these different observations together establishes that the recruitment efficacy of lateral connectivity evoking Surround responses, hence their recording detectability, strongly depends on the level of spatial summation, and is selective to the orientation feature of the stimulus flashed in the periphery (26, 27, 45).

#### 1.2. Latency dependency of Surround responses with relative eccentricity

In addition to the spatial extent of the RF, a pivotal observation, on which our working hypothesis is grounded, is that the further away from the MDF center, the latency of subthreshold surround responses progressively increases in a linear fashion with relative eccentricity. Fig. 2A illustrates the spatio-temporal latency maps of depolarizing and spiking responses of a V1 cell for sparse noise (top panel) and of synaptic subthreshold PSPs for optimally oriented long bars flashed at different eccentricities across the RF width axis (bottom panel). In this example, because spatial summation is more efficient for optimal lines than squares, surround responses become detectable on a much larger spatial extent. By measuring the onset response latencies at each tested eccentricity in the 1D-plots of ON- and OFF-responses, and applying linear regressions between the paired sets of values, one can infer estimates of the apparent speed of horizontal propagation (ASHP), ranging between 0.18 and 0.38 mm/ms. A similar observation can be done using optimally oriented and phased isotropic annular gratings centered on the RF. In the example illustrated in Figure 2B, spatial summation and response detectability still increase and the resulting latency shifts can reach up to 20-50 ms for relative eccentricity values as large as 11°.

On the whole, by comparing past and present studies using a variety of stimuli (light/dark squares, long edges, Gabor patches), we conclude that the dependency of “Surround-Only” responses with relative eccentricity from the RF center remains a hallmark of lateral processing. As shown in Figure 2C, whatever probe stimulus used, the inferred apparent speed of horizontal propagation (ASHP) remains qualitatively the same (SN in green; 1D mapping in blue, Gabor in light green, in spite of the fact that the detectability of “Surround-Only” responses depends strongly on spatial summation .

#### 1.3. Apparent Motion Axial sensitivity and Orientation tuning

Once the Gabor template optimizing the Center response was determined (see <u>Methods</u>), each RF and/or its Surround were tested with 2-stroke (Surround-Only) or 3-stroke apparent motion (AM) sequences along the cardinal RF axes (Fig. 1B). These “cardinals” axes, extending symmetrically around the RF center, correspond, one, to the “main” orientation preference axis (represented arbitrarily by the horizontal, in all the figures) and the other, to the RF width axis (vertical). For each of these axes, the orientation of the local Gabor inducer stimuli could be either the same, “**ISO-RF**”, or orthogonal, “**CROSS-RF**” to that of the RF orientation preference axis.

Examples of Surround-Only AM responses of the “cardinal” protocol are illustrated in Fig. 3A. Their magnitude can reach the size of the response to a low contrast input in the center : compare Surround-Only CP-ISO AM responses (red waveform) along the preferred orientation axis with the Center-Only response (black waveform). As a reference, the center insert in Fig. 3A illustrates the mapping of the SRF border (white contour) and the ON- and OFF-discharge fields (respectively, filled in red and blue). The surround source locations of GPs used in the AM sequence partially encroach on the SRF border, but not on the MDF. The AM evoked synaptic responses are much stronger than the impulse SN response, as shown by the absence of depolarizing responses to light (lower dark red trace) and dark (blue trace) pixel stimulations, presented in the center of the lesser eccentricity peripheral Gabor location. In this example, as in most recorded RFs, the centripetal Surround-Only AM response along the preferred orientation axis (CP-ISO, left and right red waveforms) is the strongest among the responses evoked by the Surround stimulation alone : compare it with the CP-CROSS conditions along the RF width axis (top and bottom, gold waveforms), for which responses are much weaker.

**Fig 3.**
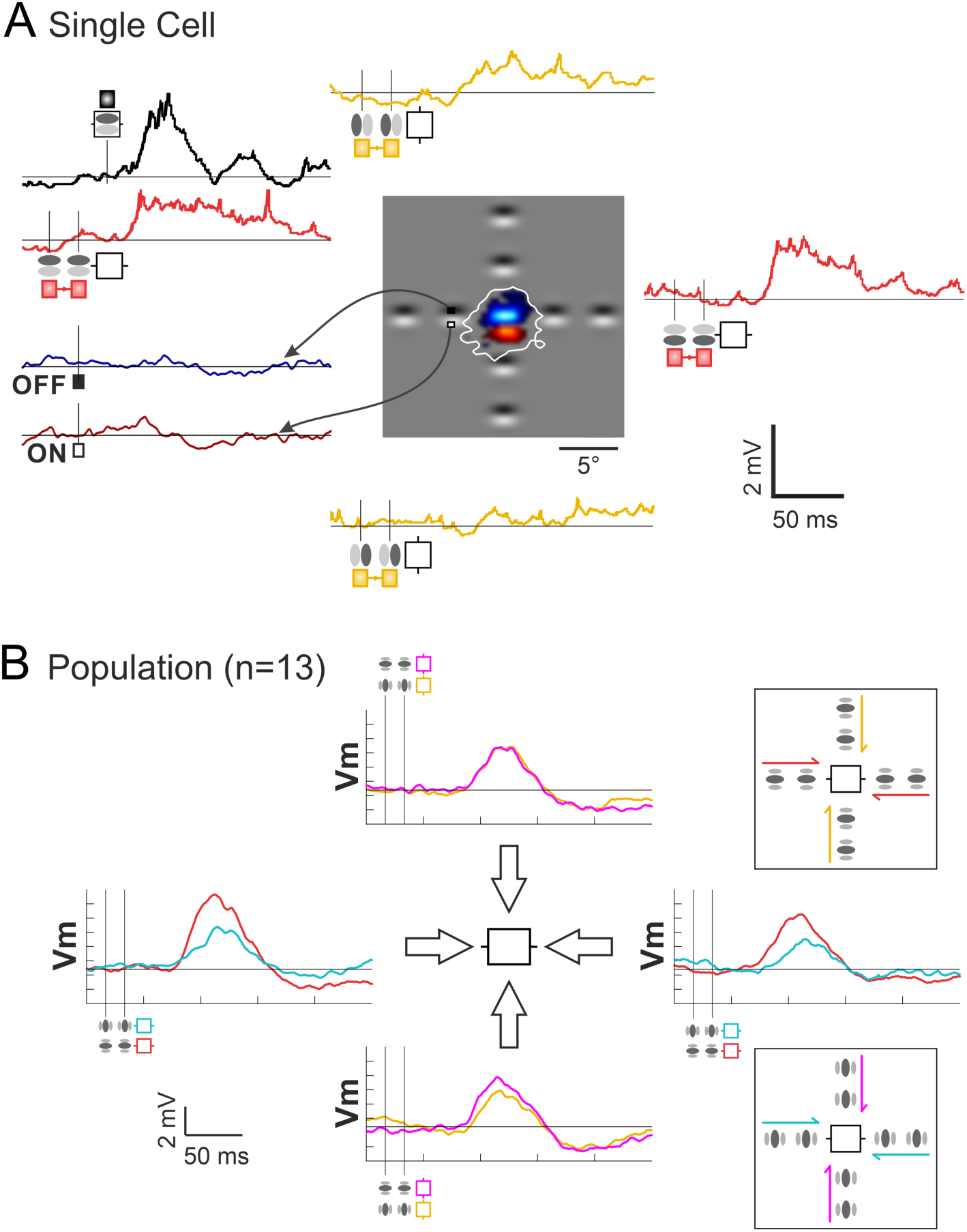
Spatio-temporal and axial selectivity of “Surround-Only” responses to Centripetal AM flow. ***[A] Single Cell example* :** Central inset cartoon : ON and OFF discharge fields, respectively in red and blue. White contour delineates the subthreshold receptive field (SRF). For comparison, respective positions of the Gabor inducer stimuli flashed in the surround are overlaid on the RF map. Two-stroke “Surround-Only” AM responses : comparison of “collinear” (red) and “parallel” (gold) responses evoked respectively along the orientation preference axis and across the width axis of the RF, with the Center-only response (black waveform). Bottom left corner, Vm subthreshold responses for sparse noise (ON (dark red) and OFF (dark blue) waveforms). ***[B] Dependency of Surround-Only responses on the orientation of the inducer*** : Right, top and bottom panels illustrate the Gabor inducer features and the color codes for the motion axis explored respectively in the ISO- and CROSS-configurations. The color code conventions are : ISO-RF: red for “collinear” along the orientation preference axis; gold for “parallel” across the RF width axis; CROSS-RF: cyan for “parallel” across the orientation preference axis; magenta for “collinear” along the RF width axis. See <u>Text</u> for details.

A complete study of the dependency of the Surround-Only AM responses on the GP inducer orientation relative to the RF preferred orientation was replicated in 13 cells. The average responses of this population have been computed by realigning arbitrarily the orientation and direction preference of each recorded cell respectively with the horizontal and right pole axes. The mean population tuning responses are illustrated in Fig. 3B, where the different AM flow conditions are schematized by color-coded arrows in the two right inserts. The top insert regroups the specific centripetal AM flows where the inducer is either “collinear” (red) or “parallel” (gold)) with the RF preferred orientation (ISO-RF condition). The bottom insert details the stimulation cases where the inducer is orthogonal with the RF preferred orientation (CROSS-RF: cyan along the main axis, magenta along the RF width axis).

This reveals two effects of the Surround contribution. The first one is a <u>non-specific</u> depolarization component found for all AM configurations, regardless of the orientation of the inducers relative to that of the RF. Globally, the recruitment of the “Near” periphery significantly rose the level of excitability of the recorded cells (Wilcoxon paired signed rank test on integral activity values between the condition of interest and the resting state, n=13, p<0.001). At the single cell level, a significant depolarization compared to the resting state (one sided z-score, p<0.05) was evoked in 76% of cells for the CP-ISO “collinear” (red) condition, and in 55% in the “parallel” (gold) condition at high AM speed. Pooling together, for each cell, responses to all CP configurations, significant spiking responses were observed in 59% of the cases. Since the classical MDF center location was kept unstimulated, the most likely explanation is that, in the “cardinal” protocol, all centripetal Surround-Only flows still recruit synaptic input originating from the feedforward SRF in addition to the proximal horizontal input.

The second component of the surround contribution is <u>orientation selective</u> and its impact can be seen in Figure 3B, riding on the top of the non-specific component. It reflects the dominance of Surround responses evoked by centripetal flows of Gabor elements “co-aligned” with the RF preferred orientation (horizontal axis in Fig. 3B) : the evoked depolarizing Surround-Only subthreshold response elicited by CP-ISO collinear flow along the main RF axis (red trace, horizontal axis in all figures) is, on average (n=13), 2.6 times larger than that evoked by CP-ISO- (magenta) or CP-CROSS- (gold) flows along the RF width axis, and that evoked by CP-CROSS-flows (cyan) across the RF main axis. A strong axial response heterogeneity is indeed noticeable in the polar representation of figure 3B, when considering separately the main and the RF width axes. On the one hand, the comparison of the CP-ISO and CP-CROSS mean subthreshold response profiles shows a clear orientation and axial bias of the “Surround-only” responses for Gabors co-aligned along the orientation preference axis (red waveforms). On the other hand, an absence of local feature selectivity was found for AM along the RF width axis (magenta and gold traces).

Globally, the comparative analysis between each configuration of Surround-Only stimulation validates the prediction of the “Dynamic Association Field” hypothesis, a concept defined in our previous study (29): the collinear configuration promotes input collection along the orientation preference axis, resulting in the emergence of centripetal axial direction selectivity for high-speed co-aligned stimuli.

### 2. Specificity of the interaction between “local” (inducer) and “global” (AM motion) features

#### 2.1. Inducer co-alignment with the RF orientation promotes centripetal synaptic integration along the global motion axis

The following contextual analysis focuses now on the modulatory effect that a two-stroke activation of the silent Surround (along each of the cardinal axes) can produce on the response to a subsequent test Gabor stimulus (the third stroke), flashed at the preferred orientation within the MDF (right column in Fig. 1B). In addition to stimulations restricted to the Surround, the contextual impact of the Surround was measured (in an interleaved way) by comparing the response to the full apparent motion sequence terminating in, or departing from the RF (3-stroke AM, <u>centripetal</u> (CP) “Surround-then-Center” and centrifugal (CF) “Center-Then-Surround” conditions) to the response to the test stimulus in the RF (“Center-Only” condition).

The complete set of CP- and CF- configurations, with Gabor inducers either co-aligned with the motion axis along the RF preferred orientation (“collinear”), or orthogonal to the RF width axis (“parallel”), was tested in 23 cells. As expected from the study of Surround-Only responses (section 1.3), both centripetal “collinear” and “parallel” centripetal AM flows invading the RF elicited a significant facilitating effect on the Center test response (WPSR, n=23, 2 conditions: AM Center+Surround vs. Center-only; p<0.001) both at the subthreshold (Vm) and spiking levels. Fig. 4 illustrates the case of a highly selective dependency of contextual control on the flow direction, emphasizing the functional facilitatory impact of centripetal collinear (CP-ISO) flows (red profile, upper panel): in addition to the change in response integral, a significant shortening is observed both at the subthreshold (upper waveforms) and spiking (lower filled PSTHs) levels (compare with black profiles of the Center-Only response) (WPSR test, n=23, 2 conditions, p<0.01). In contrast, a late change in the PSTW waveform integral, but not of its latency, is observed when the AM flow is centrifugal (CF) and proceeds from Center to Surround (green and brown, bottom panel of Fig. 4).

**Fig. 4:**
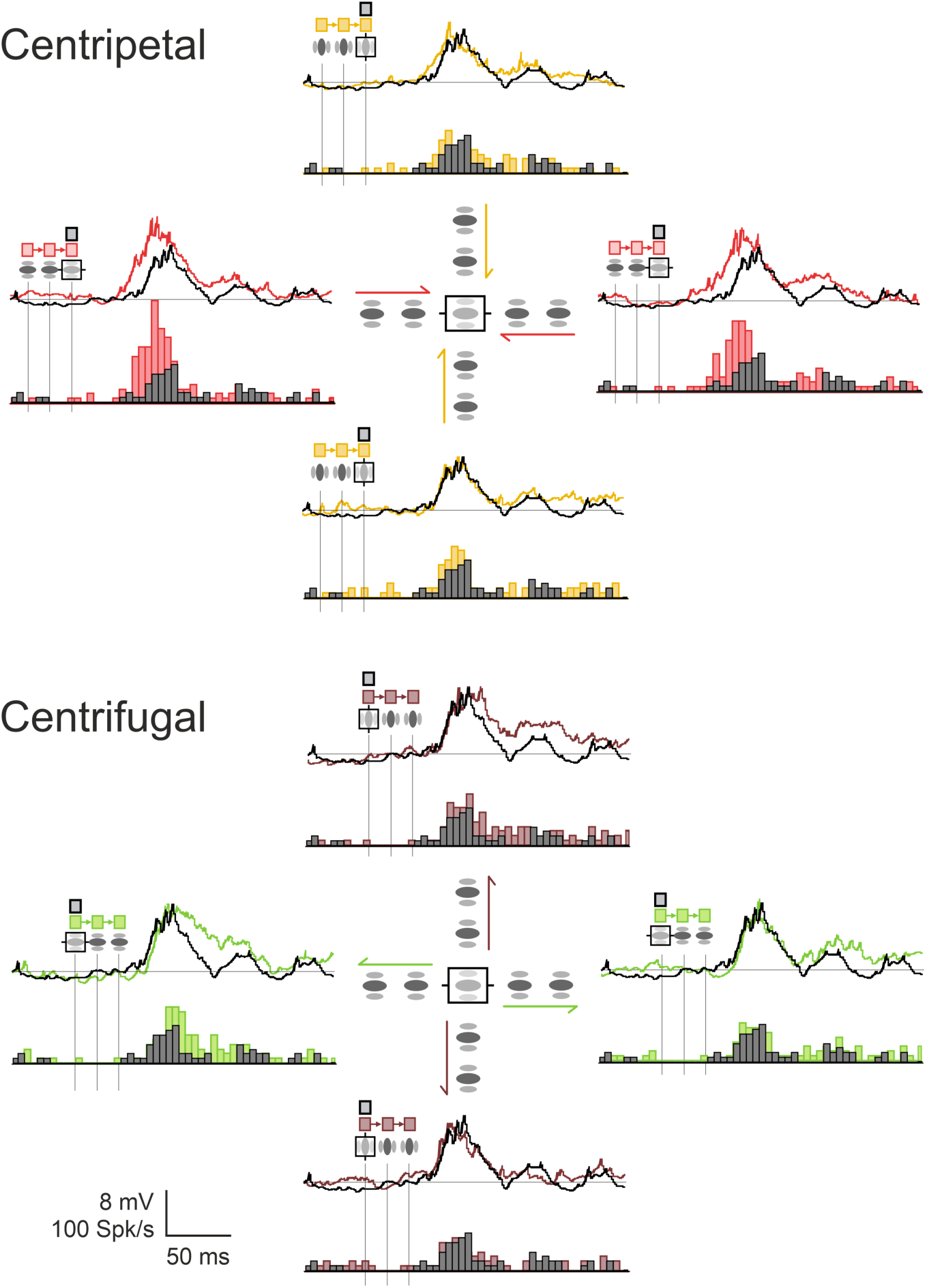
Example of contextual modulation of the Center-response by Centripetal and Centrifugal AM (“cardinal” protocol). Example of Center-Surround AM responses, compared to the Center-Only condition. The comparison, in the same cell, between centripetal (CP: top panel) and centrifugal (CF: bottom panel) motion flows illustrates the importance of the Surround-then-Center timing in the associative effect. For each configuration, a polar representation of PSTWs (top) and PSTHs (bottom) is shown for the four AM axes. In each central inset, the relative positions of the GP inducers are indicated in relation with the receptive field (rectangle, horizontal for the preferred orientation. The contextual responses are overlaid on the Center-Only stimulation (black). Note that the contextual latency shortening and the spike discharge rate increase of the Center response are observed only in the collinear Centripetal condition (top, red). Color code conventions for AM flow (arrow): CP-ISO-RF: red for centripetal “collinear” along the orientation preference axis; CP-CROSS-RF: gold for centripetal “parallel” across the RF width axis; CF-ISO-RF: green for centrifugal “collinear” along the RF main axis, CF-CROSS-RF: brown for centrifugal “parallel” across the RF width axis . See <u>Text</u> for details.

This example illustrates our main findings at the population level : First, at the Vm level, a stronger facilitatory impact was found for centripetal (CP) over centrifugal (CF) sequences (WPSR, n=23, 2 conditions, p<0.005). By calculating the ratio between the “significant” response integral (one sided Z-score, p<0.05) of centripetal vs centrigugal AM sequences, the CP/CF bias in responsiveness had a median value close to 2.0 (median ± MAD: 1.82 ± 0.42<u>). Second, the centripetal</u> modulatory effect of the test response was also selective of the motion axis and was stronger for collinear activation along the RF main axis (ISO-RF) than along the width axis (CROSS-RF) (WPSR, n=23; 2 conditions for each cell ; p<0.001). By calculating the ratio between the “significant” response integral (one sided Z-score, p<0.05) of CP AM sequences and the Center-Only test response, a facilitation by the Surround of the test subthreshold (Vm) response was observed respectively in 95% of occurrences for “collinear” and 69% for “parallel” conditions. A similar finding was also found at the spiking level, with respectively 84% of facilitatory effects for “collinear” and 66% for “parallel” conditions. The ISO-CROSS feature bias in responsiveness (median ± MAD) was respectively equal to 1.45 ± 0.28 for Vm and 1.36 ± 0.37 for the spiking activity. Finally, collinear centripetal AM sequences was found to be the only condition which induced a significant positive phase advance compared to the test response (paired statistics at the Vm level, WPSR test, n=23, 2 conditions, p<0.01).

These differential contextual kinetic changes, modulating both response integral and PSP onset latency, fit qualitatively with those expected from a simple summation model, recapitulating the respective timing in the recruitment order of feedforward and lateral inputs : CP-AM sequences tend to impact the early phase of the response when the stimulation of the Surround precedes that of the Center, whereas CF-AM sequences tend to affect the late phase of the response when stimulation of the Center precedes that of the Surround.

#### 2.2. Differential impact of “Near” and “Far” Surround recruitment

In order to better delineate the contribution of the “Far” periphery, we designed a more conservative stimulation protocol (the “radial” protocol, Fig. 1C) where the Surround stimulation was allowed only outside the impulse SRF (and not only the MDF, as done previously in the “cardinal” protocol). The spacing between GP inducers was increased from 120% of the MDF to 100% of the SRF. The absence of overlap between the nearest Surround stimulus and the SRF was checked for each of the recorded cells. Stimulation was no longer restricted to the main and cardinal axes but extended to angular SECTORS surrounding the original cardinal axes by two neighboring (± 30°) stimulation axes, so as to match the spatial sensitivity of the synaptic association field (29). We also dropped, from the stimulation set, the AM configurations used in the “cardinal” protocol for which non-specific component responses were evoked (when co-varying the local inducer orientation and the motion axis).

This enabled us to focus on our main conditions of interest with GP elements ISO-oriented with the motion axis always presented along the RF preferred orientation axis (CP-ISO and CF-ISO). Elements cross-oriented to the motion axis only referred to stimulations along the width axis of each recorded cell (CP-CROSS). Note here the terminology change in ISO and CROSS conditions - imposed by the radial symmetry – from that used in the formal (“cardinal”) protocol, where collinearity and cross-orientation the ISO-were defined relative to the RF preferred orientation axis (and not the motion axis). In addition, in the “radial” protocol, each of the 37 cells recorded was stimulated with an extended benchmark of contextual 5- to 6-stroke AM flows, richer than in the “cardinal” protocol. The parametric systematization of the various flows of ISO-axis or CROSS-axis GP sequences is detailed in each row of Fig. 1C.

Fig. 5 illustrates important spatial differences in the Surround periphery recruitment between the two protocols (“cardinal” in A; “radial” in B), by comparing the respective time-course profiles of the intracellular responses, evoked by the sole stimulation of the surround (Surround-Only, dotted traces) and by the complete AM sequence terminating in the RF center (Surround-then-Center, continuous traces). The top row in Fig. 5 illustrates the RF maps of the spiking discharge fields (ON (red) and OFF (blue)) and of the subthreshold SRF (topological union of ON and OFF depolarizing fields) delimited by a white contour. In order to focus the comparison on the stimulation axes used in the “cardinal” protocol, we illustrate, for the “radial” protocol cell (B, right panel), only the CP-ISO and CP-CROSS Sector AM configurations (first and third row, in the left column of Fig. 1C, red and gold schemes) along respectively the RF main and width axes. A straightforward observation directly emerges from the comparison of the visual field paving with Surround GPs applied in each protocol : for the same spatial viewing field size centered on each RF (20°x20°), the two Surround sources of stimulation are visible for the “cardinal” protocol, while only one - the most proximal (to the RF center) - appears for the “radial” protocol. The closest GP location in the periphery (D1 position in Fig. 1A) partially encroaches on the border of the SRF in the “cardinal” protocol, while this is not the case (by design) for any surround sources in the “radial” protocol.

**Fig. 5:**
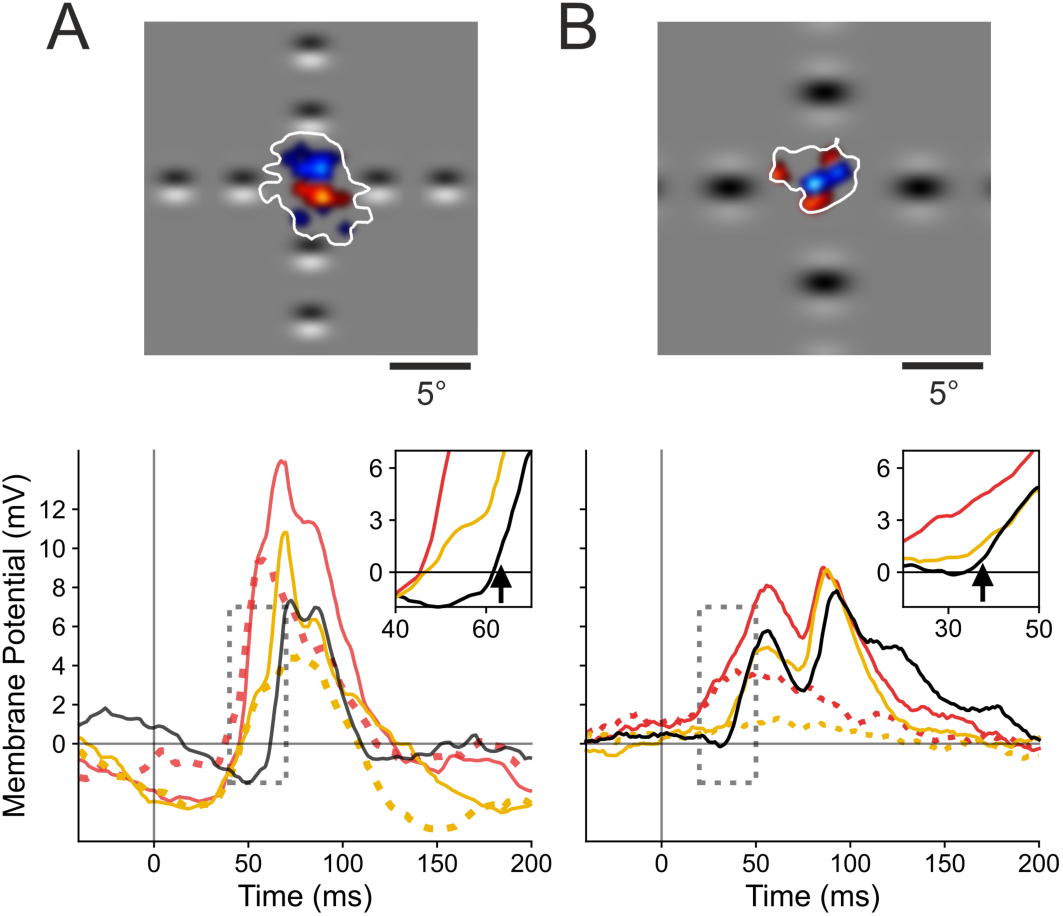
Differential recruitment of “Near” vs “Far” Surround between the “cardinal” vs. “radial” protocols. **[A]: *“cardinal” protocol*** (left panel), and **[B]: “radial” protocol** (right panel). Top row, receptive field maps. Same convention as in Fig. 3A. Only the Surround GPs on the RF main (horizontal) and width (vertical) axis (common to both protocols) are represented within the same viewing field (20°x20°). Note the encroachment of the most proximal GPs in the “cardinal” protocol over the SRF (white contour), whereas no spatial overlap is seen in the “radial” protocol. Bottom rows, the CP-ISO (red) and CP-CROSS (gold waveforms) are overlaid for the “Surround-Only” (dotted) and the “Surround-then-Center” (continuous trace) conditions and compared to the “Center-Only” test condition (black). Insets show respectively an expanded version of the PSPs onset on a 30 ms time window (dotted grey rectangle). For each protocol, arrows indicate the change in PSP slope due to the feedforward synaptic drive triggered by the Center test stimulus in the D0 location. Note that the CP-ISO (red) and CP-CROSS (gold) response latencies are the same for the “cardinal” protocol, suggestive of a common non-specific input. In contrast, the phase advance is seen only for the CP-ISO configuration in the “radial” protocol. See <u>Text</u> for details.

This difference in the spatial overlap between the most proximal “surround” stimuli and the SRF accounts for the differences in intracellular response magnitudes observed in each protocol. In the “cardinal” case (Fig. 5A), the “Near” Surround-Only stimulation evokes sizeable depolarizing responses, both for CP-ISO and CP-CROSS stimulations (dotted red and gold waveforms). Both types of CP Surround-then-Center AM flows strongly amplify the evoked PSP amplitude, although the peak amplitude gain is the largest in the CP-ISO condition. In contrast, the typical behaviour observed in the“radial” protocol (Fig. 5B) shows a different profile. For a comparable magnitude of the Center-only response (8.0 mV (“radial”) vs. 7.4 mv (“cardinal”)), the purely modulatory influence of the CP-ISO AM flow restricted to the “Far” Surround was generally much weaker (2-3 times) than that seen for the “cardinal” protocol (here, 3.9 mV against 9.8 mV). In contrast to the CP-ISO condition, the CP-CROSS Surround-Only stimulation almost did not induce a contextual response (compare the dotted red and gold traces in the bottom right panel in Fig. 5B). As a consequence, in the “radial” protocol, the complete CP-ISO AM sequence (terminating in the RF center) is the only configuration inducing a phase advance and a significant amplification of the early Center-only response.

A mechanistic difference between the two protocols is further attested by the comparison of the rising slopes of the time-profiles of the AM responses with that of the feedforward response (Center-Only). In the “cardinal” example cell (bottom left panel in Fig. 5A), both the CP-ISO Surround-Only and the CP-ISO Surround-then-Center evoked PSPs have the same initial rising slope as the Center-Only response. The latencies, when corrected for the 16 ms ISI interval, are the same in D1 and D0, thus suggestive of a direct synaptic feedforward input signature even for the D1 position (thus overlapping with the RF). Furthermore, both the CP-ISO and CP-CROSS synaptic responses start to rise simultaneously in the “cardinal” case, which indicates that the “Near” stimulated periphery recruits feedforward synaptic drive (common to D0 and D1). In contrast, in the case of the “radial” example (bottom right inset in Fig. 5B), the early phase of the CP-CROSS waveform is indistinguishable from that of the Center-Only (gold and black) waveform. The initial rising slope of the CP-ISO response (red) occurs much earlier (15-20 ms ahead) and is initially much slower than the fast rise seen for the Center-Only response (black). These latter features are suggestive of a graded spatio-temporal diffusion process compatible with horizontal activity propagation. This qualitative comparison clearly emphasizes that, by preventing the closest surround-location from encroaching on the SRF border, the non-specific tonic component of the boosting of cortical responsiveness seen in the “cardinal” protocol is no longer recruited in the “radial” protocol.

Remarkably enough, on the whole population, the “radial” protocol led to a drastic decrease of the proportion of cells showing “Surround-Only” responses (32 % (“radial”: point by point permutation test regarding spontaneous activity for a minimum of 15 consecutive ms, 10^4^ repetitions, p<0.01)), vs. 100 % (“cardinal”: Z-test above background activity, p < 0.05)). Compared to the “cardinal” protocol, it also produced weaker Surround responses, that never reached the peak amplitude of the Center-Only test response, and, accordingly, to a smaller – although significant – facilitatory modulation of the Center-response. We conclude that, in contrast to the “cardinal” protocol, the “radial” stimulation of the Surround no longer recruits as efficiently the “Near” periphery, which seems to integrate both lateral and feedforward input. Consequently, the “radial” protocol appears best tailored to detect the selective contribution of purely lateral recruitment of the “Far” periphery.

#### 2.3. Anticipatory “filling-in” responses (“radial” protocol)

To highlight the emergence of a lateral anticipatory wave due the sole recruitment of the distal periphery (Surround-Only wave) and quantify its functional role in the contextual gain control of the response evoked by the feedforward drive, we determined - in the “radial” protocol - the individual significance of each cell response to the Surround-Only condition, using a point-by-point randomization test (10^4^ repetitions, p < 0.01). The largest proportion of cells showing significant Surround-Only responses was found for the CP-ISO condition (12 cells (32% of the population)). A single cell example is illustrated in the left column of Fig. 6. A complete CP-ISO sequence invading the RF Center reveals a clear boosting effect (compare continuous red (AM) and black (Center-Only) traces, top and bottom panels). In spite of the fact that the closest surround location was sparing the SN-mapped SRF, a sizeable significant change in membrane potential was still observed in the AM sequence restricted to the surround, from D5 to D1 locations, in the absence of D0 stimulation (bottom left graph in Fig. 6, dashed red trace). For this cell, the combined dynamic recruitment of the GP sources along the AM motion axis shows a progressive build-up signal which anticipates, by ten milliseconds, the onset of the feedforward signal evoked by the Center-stimulus. In contrast, the amplitude of the linear predictor waveform - based on individual static responses to GPs in isolation – remains close to the resting state, even at the onset of the Center-Only response (green curve, bottom left panel in Figure 6). Note that the Surround-Only trace (in red) trespasses the upper limit of the confidence interval (shaded green envelope) tested by point by point permutation between the Surround Linear Predictor (SLP) and the actually recorded response to the Surround-Only sequence (10^4^ repetitions, (p<0.01)). Thus, in this particular single cell example, the AM sequence recruits a non-linear summation process of subliminal surround sources. The significant deviation from the linear prediction attests for the role of spatio-temporal synergy in maximizing the functional impact of horizontal propagation.

**Fig. 6:**
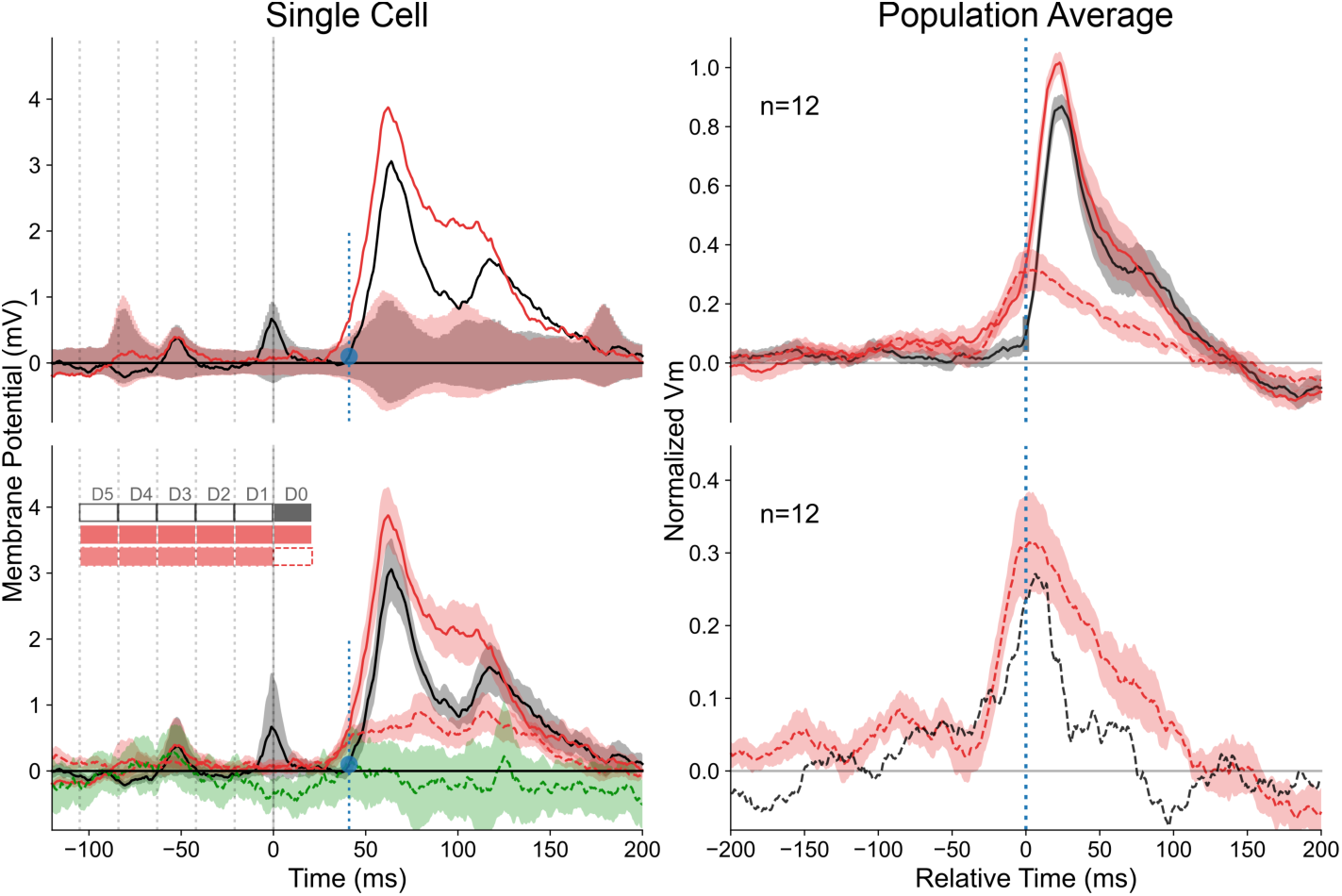
“Filling-in” responses evoked by Surround-Only AM (“radial” protocol). The analysis here is restricted to cells where Surround-Only CP-ISO stimulation evoked a significant response (n=12, see <u>Text</u> for criteria). Trace color code : black for Center-only; red for Surround-then-Center, dashed red for Surround-Only, dotted green for Surround Linear Predictor (SLP). The left middle horizontal insets respectively show the chronograms of the stroke-by-stroke stimulation sequences for Center-Only (1 stroke, filled black box, upper line), Surround-Then-Center (6 strokes, filled red, middle line) and Surround-Only (5 strokes, filled pale red, bottom line) protocols. Empty boxes indicate omitted stimuli. The time onset of each GP stroke is labeled by a thin dotted vertical line. In this figure and the following ones, the blue dot and vertical dotted blue bar on the y-axis indicate the “threshold” amplitude change of the Center-Only control curve from rest, above which statistical significance of the response is reached (p<0.01) for each individual cells. Their abscissa serves as a reference for the temporal realignment of the different contextual responses. **Left** : Single cell example. **<u>Top left</u>**: The confidence intervals (permutation test; 10^4^ repetitions, p<0.01) for Center-Only and Surround-then-Center compared to Blank are represented respectively by gray and pale red envelopes. The complete CP-ISO AM sequence (D5 to D0) evokes a significant facilitation of the Center-Only Vm response. **<u>Bottom Left</u>** : The red and black curves represent the response averages across trials (shaded area: ± SEM). When omitting the D0 stroke, the recruitment by the CP-ISO AM flow, although limited to the silent surround (D5 to D1), still induces a significant depolarizing activation (Surround-Only : dashed red). The temporal profile of the lateral wave of activity matches the “predicted” invasion of the RF Center (black), had it been stimulated. The build-up of the Surround response during AM departs significantly from the sum of the static responses evoked by each distal GP stroke in isolation (Surround linear predictor (SLP) : dashed green, confidence interval of a significant difference between SLP (predicted) and Surround-Only (observed) waveform, p<0.05). **Right, top panel** : Average response profiles for Surround-then-Center, Center-Only and Surround-Only conditions (n=12). **Right, bottom panel** : the CP-ISO Surround-Only response (mean: dotted red ± SEM: shaded area) is compared (with a different ordinate scale than in the top right panel) with the “expectation” (dashed black trace) obtained by subtracting the Center-Only (black trace, top right) from the complete CP-ISO AM response (red trace, Top right).

In the subpopulation of cells showing significant CP-ISO Surround-Only responses (point by point permutation test, 10^4^ repetitions, p < 0.01), paired comparison between responses to Surround-only sequences and their Surround linear predictor revealed that 6/12 cells (50% of significant cases; 16% of the entire population (n=37)) showed significant anticipatory responses in the CP-ISO condition larger than that predicted by the linear summation of individual GP responses. For the remaining cells, observed responses were indistinguishable from their Surround linear predictor. Note also that the slight difference between the dotted black and red curves in the Figure 6 right bottom panel – potentially indicative of a sublinear center/surround summation - could simply be due to a shift in reversal potential during depolarization.

When pooling altogether Surround-Only significant cases and realigning individual cell responses on the onset time of their respective Center-Only responses, the population average shows a mean depolarizing wave riding clearly above the noise level. This wave, depicted in the top right panel of Fig. 6, peaks at an average value corresponding to a third of the Center-Only test response peak amplitude. It corresponds to absolute values ranging between 1 to 4 mV (0.35 of normalized amplitude for the dotted red average trace vs 0.85 for the black trace). Remarkably, for the optimized AM speed, the Surround-Only average response reaches its maximum at the time of expectation of the Center-Only response. These findings strongly suggest a causal relationship between purely contextual lateral presynaptic information and its postsynaptic integration during complete apparent motion terminating in the RF center. This observation does not reflect simply a trivial non-specific summation effect : i) the gain control effect is non-linear in half of the cases (Figure 6, left); ii) the process is shown to be selective not only of the timing of the Gabor stroke sequence (optimal speed) but also of the local feature (orientation) of the Gabor inducer (CP-ISO). In spite of similar timing sequence of presentation of Gabor patches in CP-ISO and CP-CROSS configurations, the two contextual responses are actually different.

### 3. Dependency of contextual cortical control on the spatio-temporal coherence of the AM flow

We have attempted to systematically identify and quantify the key spatial and temporal factors of the CP-ISO AM sequence which condition the effectiveness of the contextual control of the cortical gain. For this purpose, we compared for each of the 37 cells, the CP-ISO configuration with AM sequences in which the spatio-temporal structure of the GP presentation was altered (Fig. 1C). The statistical significance of the measures extracted from the mean Vm responses of individual cells (top panel in Fig. 7) was assessed by comparing each AM condition to the same reference (Center-Only). Two selection criteria were used independently, based on a significant change, either i) in the onset latency, or ii) change in response strength (integral). In addition, we also used a less stringent requirement, combining one or the other or both criteria (randomization test, 10^4^ repetitions, p < 0.05). Both congruent and complementary trends were found across the three statistical criteria we used.

**Fig. 7.**
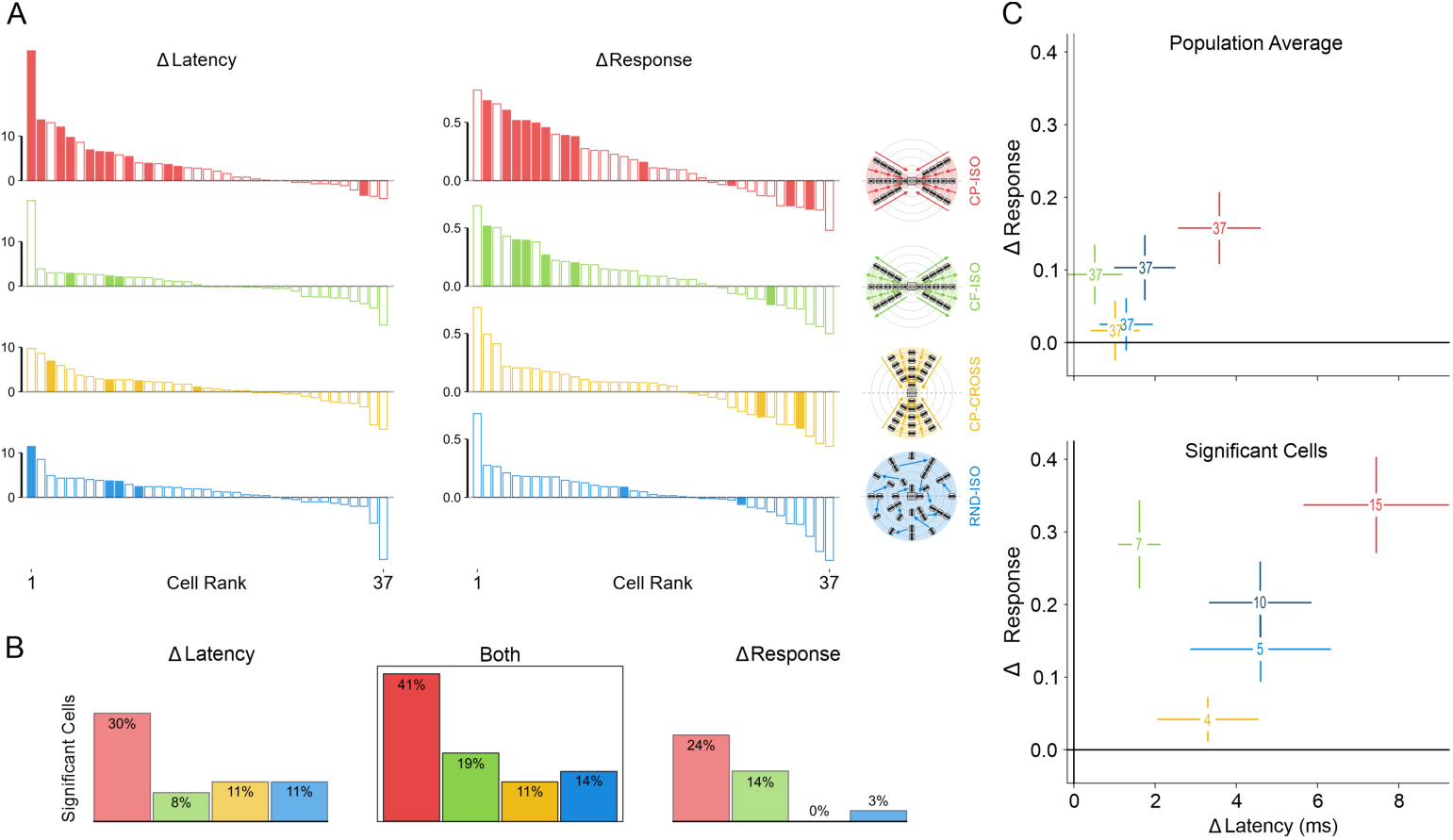
Statistical significance of contextual response changes. Color code, same as in Fig. 1C, schematized by icons in the middle column. SECTOR configuration : red for CP-ISO, green for CF-ISO, gold for CP-CROSS; dark blue for RND; FULL configuration : pale blue for RND-ISO. **A :** <u>Top left panel</u>, ranked amplitude distributions (n=37) of changes in latency (ms, left column) and in response integral value (normalized ratio, right column) of subthreshold responses (Vm), for each SECTOR AM condition of interest (first three rows). The fourth row corresponds to the FULL-RND condition (see Text for justification). Ordinates : latency shortening and integral response increase are plotted upward. Abscissa : cell rank (1 to 37) ordered independently for each AM condition. Filled bars represent “significant” individual cases (one-sided permutation test, 10^4^ repetitions, p< 0.05). Ordinates : left column, latency change (“Δ Latency”) measured at half-height of the Center-Only peak response; right column, change in the integral value of “significant” depolarizing responses (± 3σ from mean V_rest_), integrated to the point of return to baseline of the Center-only test control. The integral value change (“Δ Response“) is expressed as a normalized ratio relative to the Center-Only test response. **B :** <u>Bottom left panel,</u> each of the three histograms represents the proportions of significant cases for each AM flow condition. Left, “Δ latency” criteria. Right: “Δ Response“ criteria. Center inset (BOTH): all cases showing either a significant latency advance, or response integral increase, or both. **C,** <u>right panel</u> : Bihistogram of the Δ(response_integral) in ordinate vs. Δ(latency_change) in abscissa, averaged across conditions. Numbers and SEM correspond to values calculated across the whole population (upper plot) or restricted to the pool of “significant” cells (lower plot). Note that the global separability between CP-ISO and all other test conditions increases when restricting the pooling to the “significant” cells.

The impact of each AM configuration is expressed in figure 7A by ranked amplitude distributions of latency changes (left column) and response integral changes (right column), relative to the Center-Only condition. The proportions of significant facilitatory effects correspond to the number of occurrences where significant <u>positive</u> modulations relative to this common reference were observed. Paired comparisons between each condition (condition of interest and other controls) were also done but not illustrated for sake of clarity. The left histogram in Fig. 7B represents the proportion of “significant” latency advance cases (upwards filled bars in the left column of Fig. 7A). It is three times larger for the CP-ISO condition (30%, red column) than for the other conditions. The right histogram in Fig. 7B represents the proportion of “significant” response integral changes. The facilitation effect (positive Δ Integral) is less selective across conditions, but remains specific of the collinear configuration, since most significant cases are found in both CP-ISO and CF-ISO conditions (respectively red and green). The middle histogram inset in Fig.7B shows the impact of pooling both change criteria in a non-exclusive way (logical “OR”) : the proportion of significant facilitation cases (latency shortening OR Response Integral increase) peaks for the CP-ISO condition (41%, red histogram bar in the bottom center inset), while remaining two times larger than for the CF-ISO condition (19%, green bar) and three times larger than the other conditions (11%, gold bar for CP-CROSS; 14%, blue bar for RND-ISO). These different analyses show that the CP-ISO AM flow is by far the most effective configuration to promote the visibility of Surround-Center interactions.

For the CF-ISO condition, no consistent latency change was observed at the PSP population level (n=37), since the rising phase of the CF-induced Vm response, as expected, was generally indistinguishable from the Center-Only response. Only 8% of the cells showed a significant latency advance of a modest magnitude in the CF-ISO condition compared to the 30% found in the CP-ISO condition (green bar vs red in the left histogram, Fig. 7B). Conversely, contextual facilitation of the Center response integral was present in a slightly larger proportion of cells (14%, green bar in the right histogram of Fig. 7B). Those observations concord with those of the “cardinal” protocol, since the rising phase of the CF-induced response, as expected, was indistinguishable from the Center-Only response. On the other hand, the increase in response integral value concords with a lasting depolarization following the center-only test peak response, as more and more Gabor patches flashed at increasing eccentricity from the RF center evoke a series of lagged desynchronized lateral inputs of exponentially decreasing amplitude. The centrifugal control demonstrates that the spatial coalignment of GPs is not enough in itself to induce the observed boosting effect. Some form of spatio-temporal ordering is required such that an anticipatory flow from the periphery is generated, which, *in fine*, will boost sensory responsiveness at the time when the test stimulus hits the RF Center.

Since it is well established in most V1 cells that moving cross-oriented stimuli across the orientation preference axis of the recorded RF are ineffective in firing the cell, we focused our study of CP-CROSS AM sequences along an angular sector restricted around the width axis (Fig. 1C: left column “SECTOR”; gold condition). This choice however implies in the SECTOR configuration that the analysis relies on the comparison of response flows (CP-ISO vs. CP-CROSS) sweeping across different surround subregions, while maintaining the orientation of all GPs the same. Nevertheless, the absence of an orientation-selective component for centripetal flows of CROSS-oriented GPs along the main axis of the RF (CP-CROSS_main-axis, cyan traces in Fig. 3) justifies this simplification choice. In spite of these subtle protocol differences, only 11% of the cells showed a significant positive latency advance to the CP-CROSS stimulation, while none displayed any significant positive change in the response integral, when compared to the Center-Only test response (bottom panel, respectively left and right gold histogram bars in Fig. 7). Overall, the weak changes in latency advance or amplification of the Center-Only response found in individual cells during CP-CROSS AM flows was washed out completely when averaging across cells (RIGHT panel in Fig. 8, compare gold and black traces). A similar conclusion is reached whether the surround was partially (SECTOR) or uniformly (FULL) recruited: in both conditions (SECTOR and FULL), the contextual mean CP-CROSS responses were almost indistinguishable from the Center-Only test response.

**Fig. 8:**
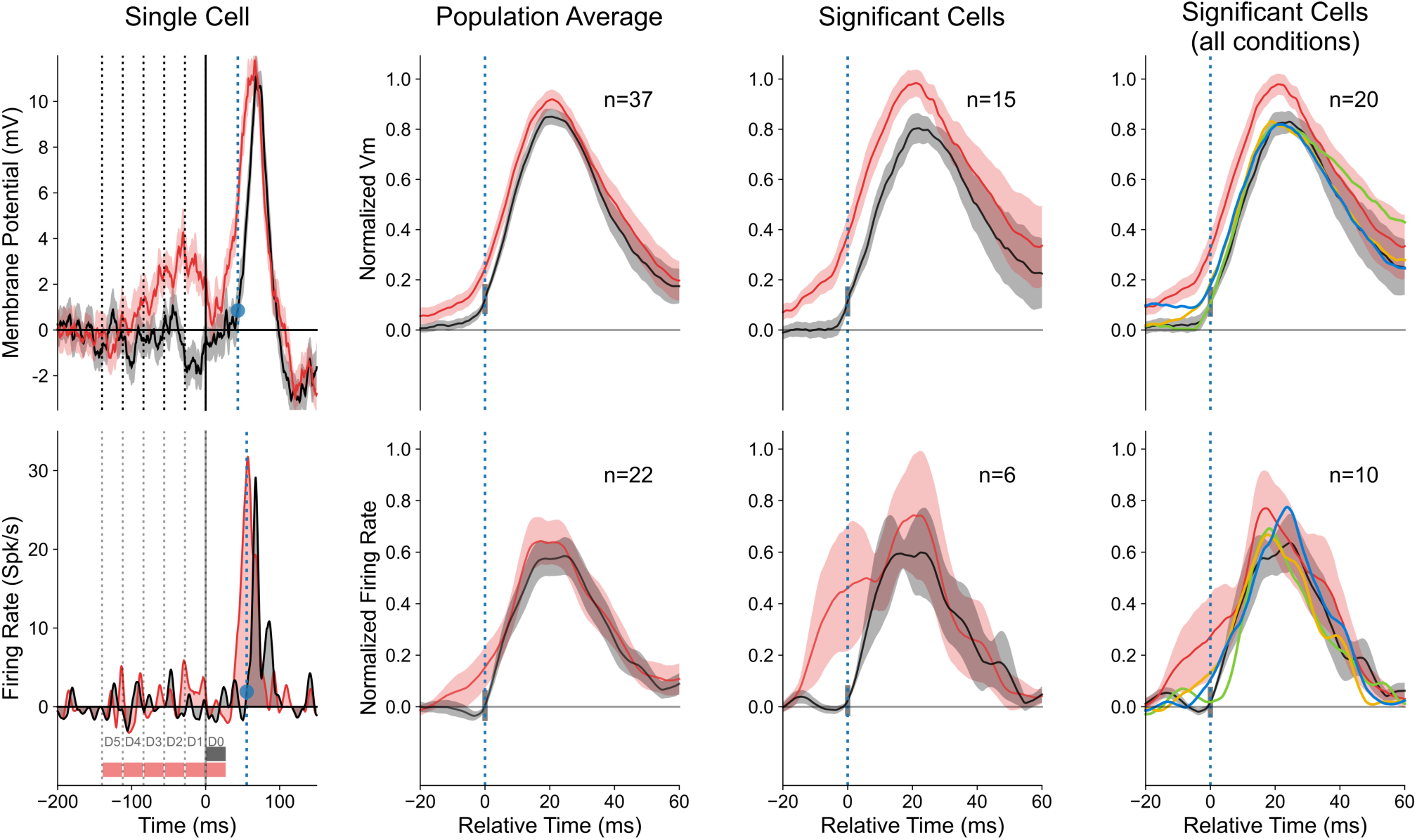
Contextual Gain Control induced by centripetal-ISO AM flow (“radial” protocol) **Top** row, subthreshold response (Vm); **Bottom** row, firing rate response (spikes/s). The first three columns, from the left illustrate the boosting effect produced by a CP-ISO AM sequence (red curve, Surround-then-Center) when compared to the response to the test GP inducer flashed in the RF Center (black curve, Center-Only). The local orientation of the GP inducers in the Surround and the global AM axis are co-aligned with the RF orientation preference. The last (right) column focuses on contextual dependency of the effect. **From left to right, first column :** Single cell example, where the vertical dotted gray lines indicate the respective timing of each peripheral stoke stimulation at successive eccentricities (from D5 to D1), whereas the continuous vertical gray line indicates the onset of the “Center-Only” test stimulation (D0). The blue dot and corresponding dotted line give the latency of the first point in time where the Center-Only response departs significantly from the resting state (p < 0.01). **Second column** from the left, population averages for the CP-ISO (red) and the Center-Only (black) conditions. Averaging is done in two steps, first across trials for each cell for the same context condition, then across cells after realigning individual mean waveforms around a common onset latency, that of their Centre-Only response (“0” mark of the “Relative Time” abscissa, blue points in left panels). Response amplitudes are normalized relatively to the peak of the “Center-Only” response. The **third column** illustrates the population average profile of individual cells showing either significant positive latency advance, response integral increase, or both, compared to the center-only test control (permutation test, 10^4^ repetitions, p<0.05). The difference in rising phases and onset latencies between the CP-ISO (red) and Center-Only (black) responses are shown on an expanded time basis. The mean ± SEM value envelopes are respectively illustrated by a continuous curve and a shaded area. **Right column**, the statistical significance analysis is extended to cells which showed significant “Surround-only” responses (subthreshold: n=20; spiking: n=10). Color code, same as in Fig. 1: SECTOR configuration, red for CP-ISO, green for CF-ISO, gold for CP-CROSS. FULL configuration, blue for CP-RND-ISO. The contextual averages show that CP-ISO (red) is the only condition where a change in onset latency is observed, both at the Vm (top) and spiking levels.

In a third control condition, the same retinal SECTOR space as in our CP-ISO condition of interest was stimulated with the same GPs (i.e., with the same number, features, and stimulus energy distribution) but the coherence of the flows was pseudo-randomized both in space and time. Such RND-ISO condition induced a slight facilitation effect on average, although significantly smaller than the one seen in the CP-ISO condition. We explored the possibility that this partial effect could be of residual nature and come from an insufficient level of randomization achievable in the SECTOR stimulation. Indeed, the number of visited nodes (i.e. GPs) is rather low (n=26), with the additional constraint that the last Gabor patch in the sequence had to be the one flashed in the Center. Note that, in contrast to coherent-AM sequences, GPs flashed during each step of the RND-ISO AM sequence could belong to any ring of eccentricity, thereby increasing at each step of the AM sequence the probability to find D1- (or D2-) to center stimulations (see <u>Methods</u>). Close examination of individual cell recordings showed that the slight facilitation most likely reflected the fact that in some RND-ISO trials, the penultimate GPs were presented occasionally by chance at the D1, or even D2 eccentricity (Fig. 1A), just one stroke ahead of the RF center stimulation. We concluded that the Sector RND-ISO sequence often recruited facilitating two-stroke AM Surround-Center pairs whose effect has already been reported (see fig. 7 in (29)).

To dilute the remnant impact of proximal-to-center interactions seen in the RND SECTOR condition, we used, as a more randomized control, the FULL stimulation configuration in which the probability to find such co-aligned D1 (or D2) GP followed by the RF center stimulation during the entire stimulation protocol was lowered. This was achieved by doubling the number of possible peripheral node locations (from n=26 to n=52), thereby increasing the level of randomization of the AM sequence. In the RND FULL mode, the stimulation was not restricted to retinal sectors on both sides of the RF but was extended isotropically to its entire Surround (bottom right blue icon in Fig. 1C). The bi-histograms in Fig. 7C illustrate the observation that the full randomization (pale blue for FULL-RND) produces changes in latency and response strength much weaker than the partial randomization limited to the SECTOR configuration (dark blue). However, still a limited fraction of the cells response to the FULL RND stimulation displayed a slight facilitatory effect (11% for latency change and 3% for response gain) compared to the Center-Only test response (blue columns, in respectively the left and right histogram bars in the bottom panel of Fig. 7B). This mild effect disappears when averaging across cells (compare average blue and black waveforms in the right column in Fig. 8). We conclude that the contextual boosting of the cortical response was largely washed out by the reduced probability of observing collinear proximal activation in the FULL RND sequences, resulting from the more extensive randomization of the flow’s spatio-temporal coherence than in the SECTOR RND condition.

Let us focus more in depth on the CP-ISO condition. The left column in Fig. 8 illustrates the typical contextual behaviour observed for Vm and spiking responses at the single cell level for the CP- ISO configuration in the “radial” protocol, when compared to the Center-Only condition : the AM CP- ISO sequence produces an anticipatory depolarizing wave that shortens the onset latencies of both the rising phases of the PSP (upper row) and the spiking response (bottom row), by respectively 14.7 and 16.3 ms. For that particular example, both a progressive build-up of a subthreshold depolarization and an anticipatory firing response are observed at early latencies, but this ramp-up feature was not systematically observed in all cells. At the population level, we found that centripetal flows composed of Gabor elements co-aligned with the global motion path along the RF main axis (CP-ISO) resulted in an overall boosting of the neural response compared to that evoked by the test Center stimulus alone.

This finding is illustrated in the second column (from the left) of Fig. 8. Note here that the PSTW and PSTHs realignments (see Fig. 6 legend) are done separately for the averaging, since a few millisecond integration lag is found between the absolute measurements of significant onset times for Vm and spikes for the Center-Only responses (see the causal shift in the vertical dotted blue lines in the single cell example). On average (n=37), the CP-ISO stimulation, in the SECTOR condition, resulted in an overall latency shortening and a slight amplification of the depolarizing subthreshold response envelope (Vm, upper row). It also led to a reduction in latency of the spiking discharge (bottom panel) in cells where spiking activity evoked in the Center-Only condition was significant (permutation test, 10^4^ repetitions, p<0.05; n=22).

In order to refine the statistical significance of the effect, we first restricted the averaging to all the cells for which, individually, the CP-ISO sequence presentation led to a significant change in the response latency and/or response integral (when compared to center-only response), whatever its sign (Figure 7A). Note here that even if the dominating effect induced by the CP-ISO stimulation was an overall clear bias toward a latency reduction, it occasionally led to a latency lengthening (negative Δ latency as illustrated in Figure 7A). We observed more globally that 1) the proportion and amplitude of positive Δ-latency and Δ-response integral was much larger than the proportion of negative ones for the CP-ISO condition of interest and 2) the proportion of significant positive modulatory effects was larger in the CP-ISO condition than in all other conditions. This led us to focus on (one-sided) <u>significant positive</u> contextual subthreshold (Vm) changes in onset latencies and/or response integral values. For this subpopulation, the CP-ISO condition produced significantly facilitated responses in 41 % of the cells (permutation test, 10^4^ repetitions, p<0.05, for 15 cells out of 37). The third column in Fig.8 illustrates the selected population average profile. Note that the quenching around the “0” time abscissa, seen for the Center-Only responses, results from the realignment process applied before averaging (see <u>Methods</u>).

To further demonstrate the contextual boosting specificity concomitantly at the synaptic and the firing levels, we tried to maximize the population size. For this last analysis, we added - to the pool of CP-ISO “significant” cells - the contingent of cells for which significant responses could be elicited by an AM flow stimulation limited to the Surround (permutation test, 10^4^ repetitions, p < 0.01; see Methods). This pooling increased the number of significant cases from 15 to 20 cells for subthreshold analysis, and from 6 to 10 for spiking analysis (one of the additional cells was not spiking). Once again, in congruence with the statistical analysis of Fig. 7, the largest proportion of significant Surround-Only responses was found for the CP-ISO condition (CP-ISO: 12/37 cells, 32%; CF-ISO: 22%; CP-CROSS; 8%; RND-FULL: 11%). The right column in Fig. 8 shows that the phase advance effect is selective of the CP-ISO condition : the response onset of the mean Vm waveform (n=20) was shortened by 8 ms and the PSP integral increased by 31%. A similar contextual phase shift is observed at the spiking level (bottom).

Altogether, our intracellular results demonstrate that a build-up of anticipatory synaptic activity is observed by sequentially recruiting local inducers co-aligned with the RF orientation axis in a centripetal way. The efficiency with which the closest surround location is recruited impacts the strength of the contextual modulation. In particular, the “Near” periphery recruitment of the “cardinal” protocol points to a bias for CP-ISO AM flows along the RF preferred orientation axis, superimposed over a non-specific facilitation component observed, at both Vm and Spiking levels for all contextual stimulation conditions. Conversely, the more precise recruitment of the “Far” Surround, achieved in the “radial” protocol, points to the selective control of the cortical gain by the CP-ISO context. This specificity truly highlights a spatio-temporal requirement in Surround recruitment in order to boost sensory responsiveness. This further emphasizes the critical need of spatial and temporal synergy between each distal surround source synaptic impact with the synaptic feedforward volley corresponding to the Center-Only drive (Fig. 1A).

### 4. Timing specificity in Predictive “filling-in” responses and Surround-then-Center interactions

#### 4.1. Synaptic mechanisms involved in predictive filling-in responses (“radial” protocol)

The central issue addressed here is to delineate how the temporal specificity of each purely contextual surround component accounts for the contextual differences in cortical gain control when lateral input is finally combined with the feedforward flow originating from the RF center. This can be studied only in cells which displayed significant responses to the sole stimulation of the Surround (point by point permutation test, 10^4^ repetitions, p<0.01). In these 12 cells, we first checked that the global contextual impact of surround and center interaction is the same as in the global population. Fig. 9 allows the straightforward comparison of the impact of the CP-ISO AM SECTOR configuration (red trace) with the three other flow configurations (light green for CF-ISO, gold for CP-CROSS and blue for FULL RND-ISO). The top left panel confirms the leftward shift advance of the Vm response by about 7 ms (top left panel, Fig. 9), selective of the CP-IS0 condition. Note that the population size is too small to warrant further statistical analysis. At the spiking level, peak values are indistinguishable, but a clear 15 ms advance only appears in spike initiation of the CP-ISO condition.

**Fig. 9:**
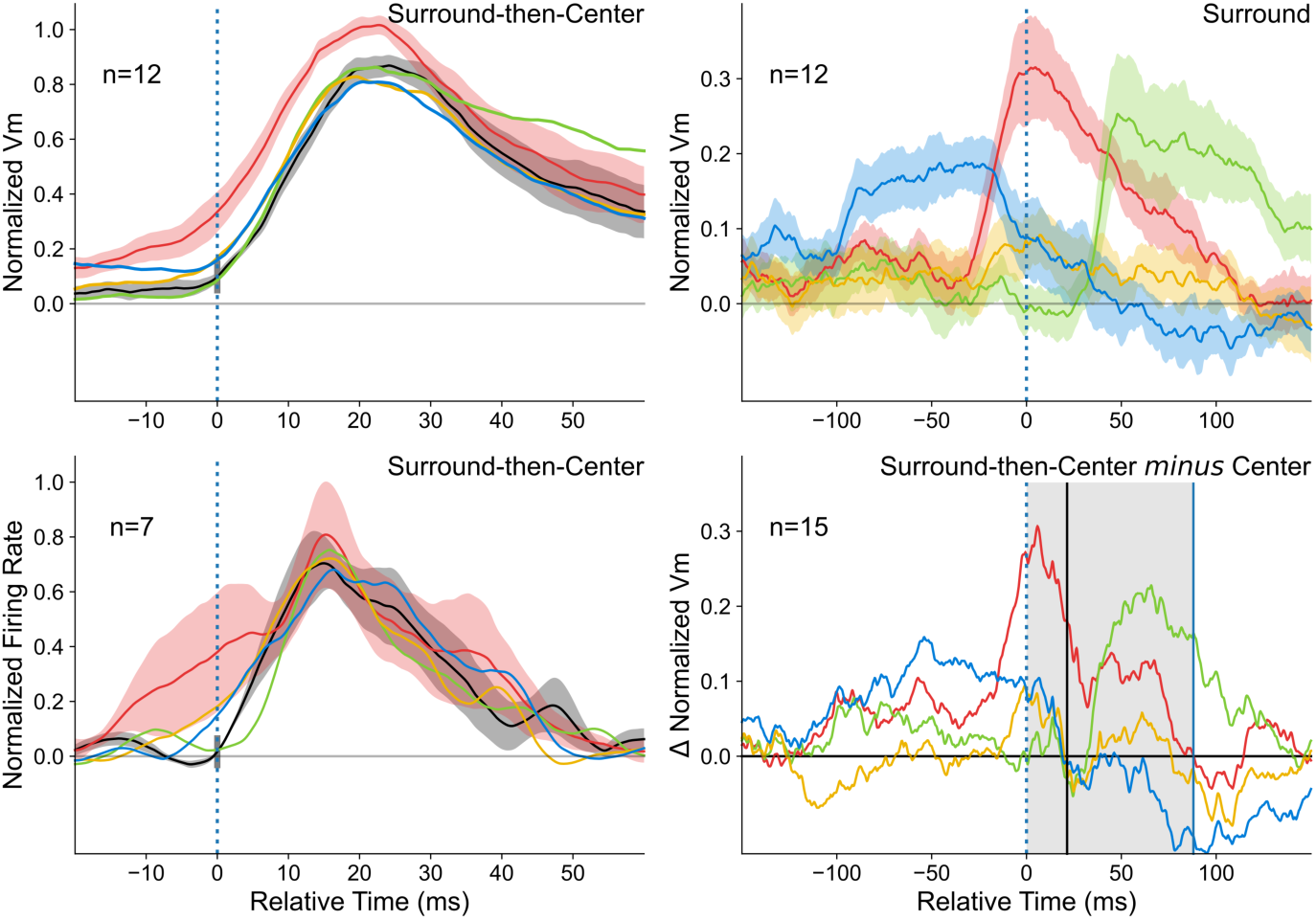
Mechanistic analysis of the “Filling-in” contribution to contextual gain control. **Left,** contextual gain control analysis, restricted to the population of cells (n=12) showing significant “filling-in” responses. Mean responses, averaged for each AM flow pattern (CP-ISO (red), CF-ISO (light green), CP-CROSS (gold) and RND-ISO (blue)) are represented at the Vm (top panel) and spiking levels (bottom) with their (shaded) S.E.M. envelopes. Same color code as in Fig. 1C. The selective latency advance and amplification of the subthreshold response induced specifically by the CP-ISO condition (top, n=12) translate at the spiking level in cells displaying spiking activity (bottom, n=7). **Right**, comparison of the contextual time course of Surround-Only responses with the predicted contribution of the Surround according to an additive model. Top, observed traces (n=12); Bottom, linear predictors of the Surround-Only responses obtained by subtracting the “Center-Only” trace from the “Surround-Then-Center” response. The predicted profiles were calculated independently for each contextual condition (same color code). The averaging process was extended to all cells showing a significant contextual change in latency and/or response integral (n=15, see Text for detailed criteria). The expected occurrence of the response evoked by the omitted Center stimulus is indicated by a grey shaded temporal window. The time abscissa of the peak of the “Center-Only” stimulus response is indicated by a thin vertical bar.

Let us focus now on the timing specificity of the Surround-Only component as a function of its mode of spatio-temporal recruitment. An elementary mechanistic approach is to compare, independently for each AM configuration (CP-ISO, CF-ISO, CROSS-ISO, RND), the temporal profile of the average Surround-only response (top right panel) to the linear prediction (“Surround-then-Center” minus “Center-Only”, bottom right panel)) obtained from the 15 cells showing significant Center-Surround modulation (top panel, third column, Figure 8). The detailed comparison of the relative time-course of “filling-in” subthreshold responses (Fig. 9, top right panel) shows that the synaptic (Vm) contribution of the “silent” periphery is the largest for the CP-ISO condition (red). Most remarkably, the Surround-only component is temporally highly selective and peaks precisely at the expected time of occurrence of the Center-Only response onset, had the RF Center been stimulated (“zero” relative time; see the dotted red trace in top right panel of Fig. 6).

This, of course, does not mean that the other contextual flows are not effective in modifying the subthreshold baseline, such as the random (RND-FULL) configuration, which shows a depolarizing plateau (blue trace) preceding the onset of the expected Center-Only response. This initial bump is compatible with the cumulative invasion (across randomized strokes) of lateral activity and corresponds to a non-null probability of stimulation by chance of the proximal periphery of the subthreshold SRF. However, in contrast to the CP-ISO AM sequence, it collapses towards the baseline before the expected onset of the Center-Only response.

For the CF-ISO condition (green trace, top right panel in Fig. 9), a mirror activation pattern regarding the Center-Only onset response is observed: starting from rest, the transient CF-ISO Surround-Only contribution growth appears only after a delay (30-35 ms), slightly after the peak of the Center-Only response had the test stimulus been presented, before displaying a progressive depolarization decay lasting up to 250 ms. This decay phase is interpreted as indicative of distal input weakening and desynchronization, the further away from the RF center (as expected from the stimulus design).

In contrast, the CP-CROSS AM sequence synaptic integration is almost ineffective (gold traces). The magnitude of its peak amplitude is several folds lower than that observed in the “cardinal” protocol, reaching a peak value of roughly 10% of the Center-Only peak response : compare gold traces in Fig. 5A, where the CP-CROSS peak magnitude of the surround-only response reaches 4.4 mV, hence 59% of the 7.4 mV center-only control peak response in the cardinal protocol vs the 1.3 mV of the Surround-Only CP-CROSS response, hence 16 % of the 8 mV of the center-only peak response. This argues once again for a “non-specific” component of the response, best seen in the “cardinal” protocol, that crucially depends on whether the most proximal peripheral Gabor patch encroaches on the SRF border or not.

An important point in this analysis is that predictors of the Surround modulatory effect, obtained by subtracting the feedforward (FF) response to the contextual response (“Surround-then-Center” *minus* “Center-Only”, shown in the bottom right panel of Figs. 6 and 9), are computed even in cells which did not show significant AM “Surround-Only” responses. In spite of the fact that the two populations used for averaging differ, a striking similarity is apparent in the overall time-courses of recorded (top) and predicted (bottom) waveforms (right panel, Fig. 9). This isomorphism globally supports qualitatively a linear (additive) synaptic integration of the dynamic AM Surround lateral contribution and the flashed Center drive..

In summary, the CP-ISO condition is the only stimulation context where the temporal profile of the Surround-Only contribution matches precisely the time-course required to interact with the feed-forward activation window (shaded area, bottom right panel in Fig. 9) and define an effective modulatory “spiking opportunity window”. In contrast, in all other flow conditions, Surround-Only contribution is out of sync with the feedforward drive and cannot exert mechanistically some form of additive gain control of the test feedforward response. Crucially, the CP-ISO exclusivity of this precise time-localized concentration in depolarizing activity power of purely contextual information actually explains the selectivity of the contextual gain during complete sequences terminating in the RF center.

Our data clearly support the selective dependency of a Surround-only activation wave on both spatial and temporal features of the AM stimulation. Co-alignment of local features in the surround with the preferred orientation of the test RF and appropriate timing in the sequential surround recruitment are both needed. If both conditions are fulfilled, the diffusion waveform, responsible for generating subthreshold synaptic responses “at the right place, at the right time”, predicts the spatio-temporal profile of the next stimulus to come (in the RF center). We therefore interpret those waves as neuronal correlates of a “filling-in” process, indicative of an anticipatory activity build-up in the cortical retinotopic region for which the “expected” visual feedforward synaptic volley is missing.

#### 4.2. Dependency of the spiking latency advance on the temporal phase between the Feedforward and Horizontal input waves (“cardinal” protocol)

The timing constraints of our protocols have a dual source: i) the synaptic activation from the surround displays a temporally constrained profile in latency basin (Fig. 2), and ii) the order of the sequential stimulation (centripetal & centrifugal) in the Surround (Fig. 4). In order to better quantify these issues, the onset latencies of the postsynaptic potentials evoked independently in the Center-Only and Surround-Only conditions were measured in the “cardinal” protocol and temporally realigned on a common reference, defined as the time at which the Center-stimulus should have been shown in the Surround-only 2-stroke AM sequence. The subtraction between the two input onset latencies will be referred in the rest of the text as the **phase** between feedforward and horizontal inputs. In spite of its weaker specificity, since the “cardinal” protocol almost systematically induced significant responses for sequences limited to the surround and strong tonic responses at both sub- and suprathreshold levels during complete sequences stimulating the center, we used the measured onset latencies of this protocol to maximize the number of cells for this analysis. This calculus was however restricted to instances where both “Surround-Only” and “Surround-Then-Center” responses were tested for each condition, in the same cells. The interest of such quantification is to give an objective measure which allows the direct comparison of the effects of different associative sequences realized for a given cell as well as across various cells (Fig. 10).

**Fig. 10:**
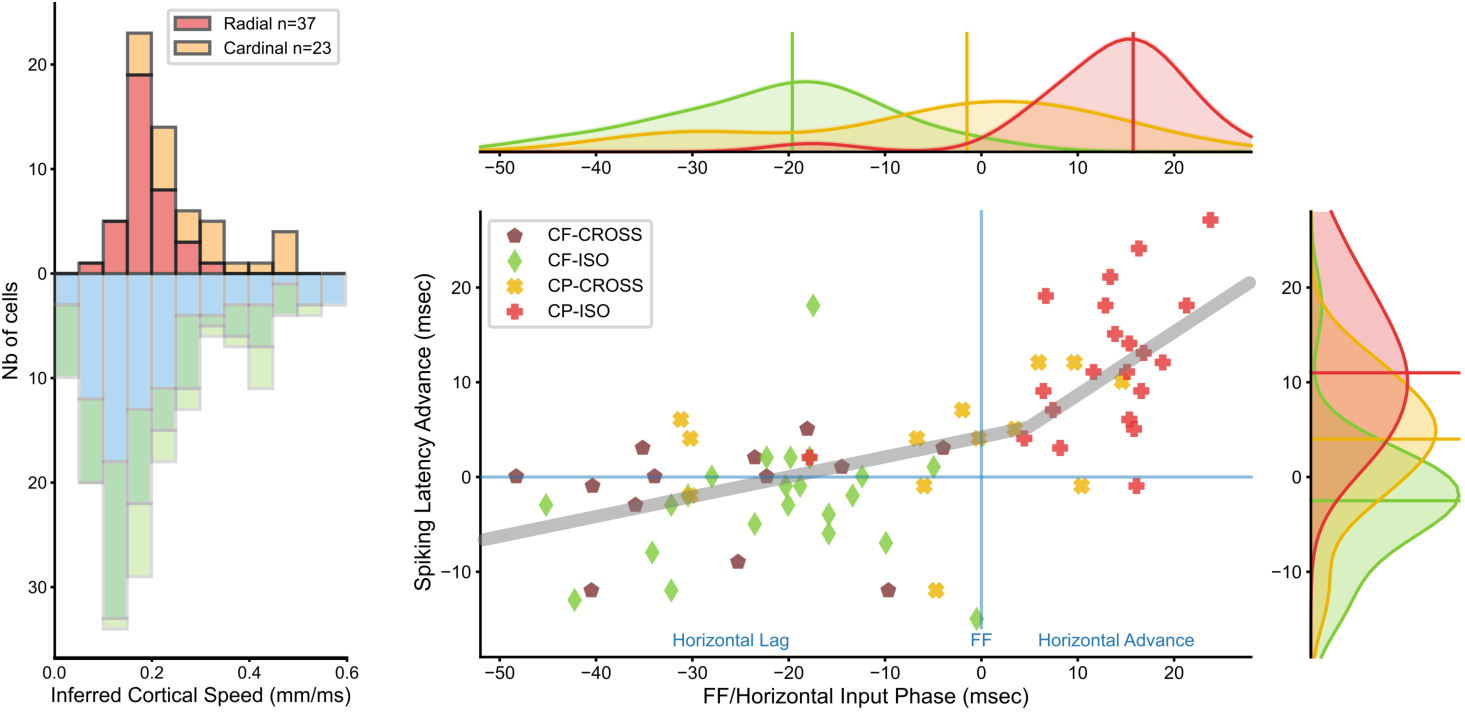
Causal impact of the temporal phase between feedforward and horizontal input on the spiking latency change (“cardinal” protocol) **Left** panel : Pooled data from the “radial” (red), and “cardinal” (orange) dynamic AM protocols (this study). Note that most apparent speed of horizontal propagation (ASHP) values, extracted from intracellular latency basin slopes with relative eccentricity, range between 0.05 and 0.60 mm/ms (with a peak between 0.10 and 0.30 mm/ms). Mirror histogram (pointing downwards) : pooled data using flashed long bars (1D-mapping, blue), sparse noise mapping (green) from (26), or Gabor patches (light green). **Right** panel: For each individual cell recorded in the “cardinal” protocol, the latencies of Center-Only and Surround-Only Vm responses are measured and subtracted, defining the temporal phase between the “horizontal” and “feedforward (FF)” inputs. The scatter plot shows the relationship between the input phase and the resulting change in spiking latency of the recorded cell produced by the AM modulation (ordinate axis). Symbols for the centripetal conditions : “x” in gold color (CP-CROSS-parallel) and “+” symbol in red color (CP-ISO-coaligned). Symbols for the centrifugal condition : “*” brown star (CF-CROSS-parallel) and green diamond (CF-CROSS-coaligned). In grey, bilinear fits. See <u>Text</u> for details. Note a significant reduction in spiking latency when the horizontal input is integrated post-synaptically ahead of the feedforward drive by 5-25 ms (phase advance, rightwards). The scatter plot data are projected on the x-axis (top row) or on the y-axis (right column) as Gaussian kernel density estimators (KDE). The color of each KDE distribution follows the convention of the “cardinal” visual stimulation protocol (Fig. 1B).

When pooling all cells (n=23 cells, 67 measures), the distribution of input phase relationships ranged from +25 ms of phase advance to -50 ms of phase lag between horizontal and feedforward input waves at the subthreshold level (abscissa in the bottom-right scatter plot of Fig. 10). As expected, a strong bias in the distribution was observed in favor of phase advance of the Surround input for the centripetal sequences (gold “x” (CP-CROSS) and red “+” (CP-ISO) symbols) and delayed activation for the centrifugal sequences (brown “*” (CF-CROSS-parallel) and green diamond (CF-ISO-coaligned) symbols). At the spiking level, the latency shortening of the contextual (Surround + Center) output, compared to the Center-Only condition, appears crucially dependent on this input phase relationship: it was only observed for centripetal condition, i.e. when the Surround response onset (the horizontal input), preceded in time the Center-only response onset (the feedforward (FF) input). The intercept of the grey bi-linear fit, shown in the scatter plot of Fig. 10, has been optimized by two half-regression lines independently of the stimulus conditions (taking all the data points together). The optimization of explained variance by a bi-linear regression composed of two connex linear segments results in a partition of the data cloud in two abscissa regions on each side of the zero phase axis: one (on the left side) where the horizontal input lags in phase the feedforward drive, and for which no significant trend is observed (r^2^=0.01; n=32, p>0.5); the other (on the right side), where the horizontal input is postsynaptically integrated ahead of the feedforward drive by 5-25 ms (r^2^= 0.356; n=37; p<10^-5^). In this latter regression domain, the observed reduction in spiking latency (latency advance pointing upwards on the y-axis, reflects in an almost linear way the phase advance of the synaptic echo of the horizontal input relative to that of the feedforward drive (“0” abscissa).

The Gaussian kernel density estimator (KDE) of the input phase distribution (along the x-axis, above the scatter plot) clearly shows that the depolarizing input wave evoked by a collinear pair of stimuli flashed in a centripetal sequence in the “Near” periphery (CP-ISO, in red) was arriving earlier (17 ms from the scatter plot projections; 9 ms for paired data with CP-CROSS, in gold) than when the stimuli were oriented orthogonally to the width axis path (CP-CROSS, in gold) (Wilcoxon WPSR test on latencies, n=39, 2 conditions, p<0.01). Note that, in some cells, the latency advance at both Vm and Spiking levels of the CP-CROSS stimulation (gold symbols) is non-null. However, the average 9 ms difference in subthreshold responses of CP-ISO and CP-CROSS conditions of the “cardinal” protocol corresponds to the 8 ms latency advance (between CP-ISO and Center-Only response), found for the CP-ISO significant subpopulation of “significant” cells in the “radial” protocol. Note that, for this precise contingent, the latency advance component of the CP-CROSS response was indistinguishable from the reference (Figs 8 and 9).

Only centripetal collinear CP-ISO sequences could induce significant shortening in output spiking latency, by an advance as large as 30 ms. As shown by the KDE distribution on the y-axis (right of the scatter plot), the median latency reduction observed for the collinear axis during a centripetal stimulation (red line on the shaded red envelope) was 11 ms. In the other conditions, either centrifugal or centripetal “parallel” along the RF width axis, no significant latency advance could be seen: the three other distribution fits are all centered on a null latency change. At the spiking level, the median difference between the onset latency of CP- and CF-conditions is of 13 ms (respectively, red and green lines).

#### 4.3. Synergy depends on the relative match between apparent motion and intracortical horizontal propagation speed

A subtle difference between the two types of protocols is that the AM speeds used in the “radial” protocol depended on the GP duration and on the size of the SRF, and not only on the size of the MDF (as this was the case for the “cardinal” case). This finer speed tuning was adjusted to the characteristics of each individual cell since the response latency and strength of evoked responses both depend on the stroke duration and the distance between strokes. For the 37 cells of the “radial” protocol, the average AM speed was of 189 ± 47 °/sec, (ranging from 72 to 312 °/s), hence below the average speed of 329 ± 101 °/s (range 175-500 °/s) used in the “cardinal” protocol. Note that despite this difference, the global ASHP distributions reported in the “radial” and “cardinal” protocols overlap closely with the overall range of distributions reported in (26) and (29) (mirror histograms in left panel of Fig. 10). A detailed comparison shows that the inferred speed of laterally conveyed activity is slightly accelerated for “dynamic” sequences of AM flows recruiting the periphery (top of the histogram) than for “static” flashed stimuli measures (bottom), peaking respectively at 0.175 (n=60) and 0.125 mm/ms (n=168). The slight shift in the ASHP distribution reflects a change in the median values (0.225 vs 0.175 mm/ms) rather than in the mean values which remain similar (0.231 and 0.251 mm/ms). Those results point to the notion that the speed of laterally conveyed activity is not completely hard-wired but depends partly on the type of visual stimulus used. Therefore, higher levels of spatio-temporal coherence along the motion path slightly increases the speed of laterally mediated activity, leading most likely to a cumulative shortening of the response onset latency at each updated locus of stimulation of the AM path in the cortex, a phenomenology already reproduced in simulations of nested horizontal propagation (47).

Functionally, a critical prediction of our “working hypothesis” is that the visibility of the lateral contextual modulation of the feedforward input should therefore increase when the speed of the AM-evoked wave in cortex grows closer to horizontal propagation speed. Focusing on the SECTOR configuration of the “radial” protocol, we studied more in depth the impact of the CP-ISO AM kinetic effects on synaptic integration in V1 in a subset of twelve cells, by comparing, in each of these cells, the impact of AM sequences of various speeds. The speed of lateral connectivity recruitment was controlled in a graded way by introducing temporal delays between GPs strokes (detailed in the chronograms in the bottom panel of Fig. 11). We first determined independently for each cell the “optimal” AM speed of surround recruitment, which set the 100% nominative reference speed value (ASHP). In addition, for each of these cells, similar AM sequences were replayed at 70%, 50% and 30% of their nominal speed value, and interleaved in a pseudo random fashion. In spite of cell-by-cell differences in their absolute optimal ASHP values (varying between 150°/s to 250°/s), population averages shown in Figure 11 were done by grouping across cells responses obtained at 100%, 70%, 50% and 30% of their respective “optimal” ASHP.

**Fig. 11:**
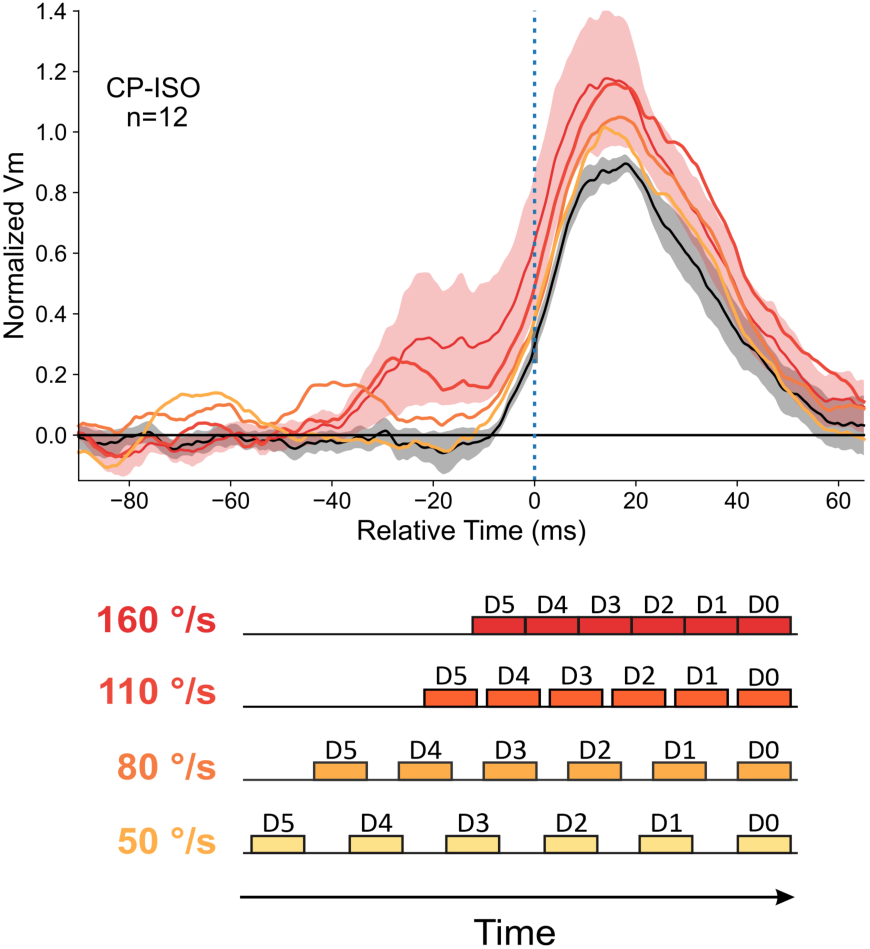
Dependency of the Contextual effect on the speed of the AM flow : **Bottom,** the chronograms depict the 4 inter-stroke interval conditions used in each cell to probe the temporal dependency of the interaction between the horizontal and the feedforward synaptic waves (color code reddens with speed value). The exact value of the “optimal” reference speed (100%) is defined on a cell-by-cell basis. Mean speed values, averaged across cells (n=12), are given left of each chronogram. **Top,** population averages for the CP-ISO SECTOR condition. All individual responses are realigned with the Center-Only response onset. The color-coded overlaid waveforms, averaged across cells, allow the comparison of the time-course of the response for the optimal speed value (red) and its S.E.M. envelope (shaded red) with that observed for various proportional reductions of the AM flow speed (from top to bottom: 70% (dark orange), 50% (orange) and 30% (light orange)). The contextual response modulation amplitude decreases proportionally to the AM speed reduction from its optimal value.

The comparison of the average contextual PSP response profiles of this subpopulation for incremental AM speeds (30%, 50%, 70% and 100% of the “optimal” value) highlights a cumulative speed-dependent anticipatory recruitment of the periphery, with a bumpy stairway-like profile preceding the main response peak (top panel in Fig. 11). Crucially, the overlay of response profiles for different speeds suggests a progressive merging of lateral inputs impinging on the central target RF, from weaker amplitude and asynchronous early responses at lower speed to larger amplitude and synchronous “in-phase”-summation with the feedforward drive (Center-Only, black curve in Fig. 11) when approaching the “optimal” AM speed. As speed value grows (from 30% (light orange) to 100% (red) of the optimal speed), lateral influence gets progressively synchronized when it reaches the RF center. This is attested by the presence of dissociated bumps (preceding the Center response) at slower speed, which likely reflects sequences of asynchronous activation by the distal surround sources. In contrast, at the “optimal” speed, the precise time-concentration increase in depolarizing activity power enables a more efficient interaction with the center-only response, thereby boosting in a causal way the feedforward PSP. Once more, this result clearly emphasizes the crucial timing of synaptic recruitment in the Surround, which controls the emergence of synergy between long-distance (orange arrows in Fig. 1A) and cascades of shorter neighbor-to-neighbor links (grey arrows in Fig. 1A) along the AM path. We conclude that “in-phase” summation of lateral inputs with the predicted time of the feedforward drive produces the shortening of latency and amplitude amplification reported during CP-ISO flow at high AM speed.

Altogether, our results show that facilitatory contextual control of V1 responses is optimized along the orientation preference axis when the global motion speed lies within the range of visual flow speed predicted from the apparent conduction velocity of horizontal connectivity. For slower motion, the horizontally-driven facilitation effect disappears and stimuli cross-oriented to the motion path, presented across the width axis, remain the best drivers of cortical cells. We conclude that the axial preference of V1 cells shifts abruptly by an angle of 90 ° when the motion speed switches from 2-20 °/s to 100-250 °/s.

In summary, the spatial congruence of the elements regarding the preferred orientation axis, the spatio-temporal coherence of the flows regarding the RF center (centripetal), the speed of recruitment of the periphery and the precise timing selectivity of Surround-only responses, all suggest that some form of peripheral subthreshold “prediction” is generated internally in V1, to interact, within the proper spatio-temporal window, with the feedforward Center-Only synaptic volley. These highly selective features justify, to our view, the qualification for the existence of neural correlate of “predictive filling-in” in V1, to be distinguished from a non-specific anticipatory depolarizing spread.

## Discussion

Most studies of contextual modulation in the early visual system are based on synchronous Surround and Center stimulation, pointing to mixed suppressive and facilitatory effects (13, 14, 48–53). We document here a dynamic regime of asynchronous Surround-Center activations, of specific speed and spatial anisotropy, which, under precise timing and co-alignment conditions, reveals a facilitatory form of contextual cortical gain control. This effect, which links global motion direction and local orientation, is strong enough in the anesthetized brain to shorten spiking response latency by a few tens of milliseconds and shift the axial preference of V1 cells by 90° for saccade-like speed, favoring contour integration along the orientation preference axis.

### 1. Orientation anisotropy, Motion streaks and Speed sensitivity

The dependency on co-alignment and motion along the RF preferred orientation axis reported here correlates with the anatomical bias for iso-oriented long range connections described in higher mammalian visual cortex (*cat*: (54–56); *tree shrew:* (57–59); *macaque* : (60, 61); *squirrel monkey* : (62)). At the functional level, it is consistent with the concepts of the perceptual “Association Field” in humans (4), the neural “facilitation field” in the behaving NHP (11, 63) and the synaptic “Dynamic Association Field” in cat V1 (29). In terms of modulatory efficiency, the requirement of maximizing the spatio-temporal synergy from the Surround corroborates the stimulus dependency in the recruitment of long-distance iso-tuned interactions reported with imaging techniques at the mesoscopic level (40).

Importantly in V1 neurons, the integration axis (the RF preferred orientation) for which the strongest responsiveness modulation by fast (150-250°/s) apparent motion is observed, is orthogonal to the preferred directional axis of V1 neurons (the RF width axis) when probed with slowly moving bars and full field gratings. Classically, area 17 (V1) receptive fields are best activated by slow motion across the RF width axis (1-2 °/s in (46); 3-10 °/s in (64), but quantitative studies reveal however a zoo of speed tuning selectivities in V1 ((65–67); review in (68)). Among these, “parallel motion direction selective” neurons could provide a potential substrate for the effects reported here. However such trigger features are classically found in higher visual cortical areas dedicated to global motion (type II MT cells (69)), but not in V1. In contrast, Geisler and colleagues hypothesized, then described, a specific class of cells in V1 responding to Gaussian blobs, named “streak” detectors, oriented in the direction of motion (70, 71). To account for disambiguation of motion direction, Geisler introduced a hypothetical multiplier circuit between direction selective and “parallel” detectors, whose existence needs still to be demonstrated (70). Realistic estimates for the speed threshold above which “motion streak” operates point to medium range values, around 20-60°/s for cat and 10-20°/s for monkey (inferred from (72)). This “streak” effect, however, depends on the test stimulus,i.e. small isotropic Gaussian blobs, critical to avoid recruiting side-inhibition, and is absent for oriented plaid patterns (71). Other studies reported related findings: Orban and colleagues showed that spot directional tuning shifted by 60-90° above a speed threshold, whereas bar tuning remained invariant (68, 73). Intrinsic optical imaging experiments - combining textures tiled with small bars and independent motion components – point to a complex dependency of orientation domains recruitment on local motif length and motion speed (74). Similarly, 90° shifts in orientation domains were described for moving dot noise above 14.5°/s in cat area 17 (75).

In contrast, the 90° shift in axial motion sensitivity reported here is found at much faster (saccade-like) AM speeds (>50°/s), and relies on the same local inducer (oriented Gabor) shifted in space and time. It is the precise time-lagged recruitment of Surround-then-Center synaptic inputs, that conditions the contextual amplification and the phase advance of the Center response. Both the binding specificity of the local (orientation) and global (motion axis) features of the AM flow, on the one hand, and the match between the optimal AM speed and ASHP values, on the other hand, support the view that the main player is long-range horizontal connectivity intrinsic to V1.

### 2. Novelty of the study

The novelty of our findings is several fold : 1) these lateral diffusion effects recorded in the anesthetized, hence non-attentive, animal are likely to be intrinsic to V1 and do not require behavioral attention and top-down control from higher cortical areas (weakened by anesthesia) ; 2) they depend on the spatiotemporal coherence of the AM sequence and grow with flow speed until matching the apparent speed of horizontal propagation (ASHP) in V1; 3) in that range of speed, they produce a shift by 90° of axial motion sensitivity, which becomes co-aligned with the preferred orientation RF axis; 4) they provide evidence for an internally-generated propagation process, binding local (orientation and position) and global (motion and direction) features; 5) they constitute a plausible neural substrate of psychophysical effects in humans reporting a perceptual bias in speed estimate for collinear motion (76).

### 2.1. Intra-V1 horizontal diffusion vs. cortico-cortical feedback

Conventional models of visual cortical processing assume that complex computations, such as motion direction disambiguation, are encoded only at higher levels in the visual cortical area hierarchy, and are absent in V1 (9, 10, 77–81)). Most electrophysiological studies in the behaving primate point to the late retroaction in V1 of a top-down signal gated by attention (82–85). They attribute the delay of several tens of milliseconds of the change in firing of V1 cells after response onset, to the late action of the cortico-cortical feedback (86–90), rather than to the slow horizontal propagation in V1. However, puzzlingly enough, axonal feedback transmission from higher areas is 10-100 times faster than intrinsic diffusion in V1 (see also (91) for the parallel recruitment of dual lateral-feedback kinetics).

The challenging view presented here is that, as early as V1, horizontal connections play already an important role in the early perception of retinal flow and global motion. The visibility and spatial spread of their functional dominance in cats & ferrets vs non-human primates (NHP) may reflect species-specific differences in retino-cortical magnification factor. Horizontal axon length (4-8 mm) is the same across species, but covers up to 8° of visual angle in cat-ferret (*<u>cat</u>* : (54, 55, 92); *<u>ferret</u>* : (59) whereas it remains within a radius of 1.5 hypercolumn (2-3°) in primates (93), which matches the spatial extent of collinear facilitation observed in human psychophysics (42, 94–96).

The multi-stroke AM protocols (“cardinal” and “radial”) were both designed to promote the in-phase synchronization of the synaptic impacts produced by the sequential stimulation of each GP flashed along the motion path with the central feedforward drive (orange arrows in Fig. 1A). Our prediction was that this synchronization would be optimized for a flow speed in the range of the apparent propagation speed of horizontal connectivity (ASHP in (26)). This view is confirmed by the speed-dependency tuning study reported here (see <u>Results</u>). However, the variability of the intracellular anticipatory Vm profile across cells suggests that other synaptic recruitment paths are likely to contribute, in particular the neighbor-to-neighbor links (grey arrows in Fig. 1A). It is thus likely that the “horizontal” drive efficiency results from a combination of the asynchronous visual activation of two classes of synaptic sources: 1) long-range monosynaptic inputs recruited at different eccentricities, whose distributed and delayed impact is integrated in phase by the target cell; 2) polysynaptic short-range input, reflecting chains of next-to-next neighbor activation (studied in our previous 2-stroke AM (29)). This second effect is amplified by the re-ignition process, imposed at each GP locus by the next feedforward input recruited on the apparent motion trajectory. Note that this rolling wave effect is not seen for single stroke stimuli, since, otherwise, the spiking discharge field of the recorded cell would extend far beyond the spatial region defined by the core thalamo-cortical projection.

Since the reverberation delay imposed by cortico-cortical loops is in the order of a few milliseconds only (97), it is plausible that most long-range interactions in the primate may operate by nested polysynaptic feedback afferents from higher cortical areas through chains of neighbor-to-neighbor lateral relays, whereas in the cat and ferret, a substantial part would remain encapsulated monosynaptically within V1 (7). Our results may provide new insights on the evolutionary complexification of cortico-cortical computation. They could also spark interest in AI and deep learning in artificial vision, where the diffusion effects of horizontal connectivity in a single V1-like terminal layer can be confronted with the backpropagation of “network belief” implemented by nested feedback connections in a multi-layer hierarchical cortical-like architecture (98).

#### 2.2. Dependency on spatiotemporal coherence and visual flow speed

Since 2-stroke AM provided evidence mostly for subthreshold synaptic amplification (29), we aimed here at promoting the efficiency of centripetal AM CP-ISO sequences to produce a spiking discharge. This was achieved by increasing the number of elementary strokes in the AM sequence, from 3 to 6 from the “cardinal” to the “radial” protocols. In the “radial” protocol, the comparison between FULL-RND, SECTOR-RND and SECTOR-CP-ISO conditions (Fig 7 right panel) provides evidence that the boosting of sensory responsiveness grows with the level of spatio-temporal coherence. The speed dependency of the effect demonstrates unambiguously that the efficiency of the contextual control of the cortical response gain and phase is optimized when a causal coherence is reached, i.e. when the horizontal wave predicts the arrival of the feedforward wave.

The optimal range of speed (150-250°/s) reported here contrasts with a recent study of anticipatory responses in the fixating monkey, best seen for stimuli cross-oriented to the motion path, in continuous slow motion across the RF width axis (6.6°-13.2°/s) (99). The authors propose a complex mechanism, based on the interplay of the “feedforward” trace imposed by the continuous motion of the stimulus and a medium range isotropic horizontal propagation wave. This activity bump still allows a suboptimal slight interaction with the feedforward drive when it propagates faster than the stimulus. Their interplay would cause a pre-activation of the cells, facilitating in turn the feedforward reactivation. This proposed mechanism is highly dependent on the retino-cortical magnification factor and does not account for the global dynamics of the anticipatory wave for distances beyond monosynaptic connection lateral extent. The propagation speed across long trajectories (> 3° from the Spiking discharge field center), well beyond the spatial extent of monosynaptic links observed in NHP (<2°), is indeed ten times slower (<= 0.02 mm/ms) than the apparent speed of horizontal propagation (0.05-0.50 mm/ms; see (26, 32)). The global kinetics are more in line with a slower “rolling wave” diffusion mechanism (see also *in vitro* (100)). The speed range of this anticipatory wave reported in awake macaque (<15°/s) is lower than in the anesthetized cat area 17 (15-40°/s in (39)), where it has been shown that a moving square is more rapidly processed than a flashed one. These different effects share some similarities with those reported here, but differ significantly in their speed range (by a factor of 10-20) and preferred axial sensitivities (CROSS-parallel vs ISO-Collinear). Furthermore, they do not require collinearity, a hallmark of horizontally-mediated V1 computation.

#### 2.3. Binding of global and local features as early as V1

The current consensus concerning cortical correlates of low-level perception mixes hierarchical processing and cortico-cortical recurrency : V1 plays a pivotal role in synchronization dynamics back and forth with high-order cortical areas (16, 101). Intra-V1 synchronization and inhibitory interactions are hypothesized to interact through a form of “competitive coding”, before the output is further processed downstream in the cortical hierarchy. A non-exclusive alternative is that reverberating activity in V1 already participates in non-attentive “pop-out” perception. We propose that horizontal connections, intrinsic to V1, are instrumental to the neural implementation of low-level perceptual Gestalt laws, in parallel or before information is broadcasted to the mosaic of higher cortical areas. Our data suggest that, in Gestalt-like configurations, horizontal connectivity propagates some kind of “network belief” (built-up on past activity ignition and shared input statistics in the Surround) to the connex V1 neighborhood, the most likely to be next stimulated (102).

Remarkably enough, the “prediction” wave, seen for “Surround-Only” AM CP-ISO sequences, binds local information acquired along the motion path. This observation, although recorded in the anesthetized brain, globally fits with Bayesian theoretical frameworks, where omission of the next-to-come stimulus in repetitive and predictive activation sequences unmasks an intracortically generated “expectation” signal mediated by horizontal connectivity. This finding has to be distinguished from the classical “predictive coding” schema (103), where what is propagated is the “residual error” from the expectation generated in higher cortical areas rather than the diffusion of the network belief context (98). At optimized speed, the self-generated “Surround-Only” wave is shown here to exhibit the same timing as the feedforward (FF) input onset evoked by the next stimulus to come, had it been presented. More exploration, outside the scope of this paper, would be needed to test how center-surround interaction is affected by the contrast (low/high) and the orientation (same/mismatch) of the test Center stimulus, and how possible sub-linearities can be interpreted in the context of the principle of redundancy reduction framework (104).

Our results suggest that V1 may solve the motion extrapolation problem just through selective internal diffusion. Our interpretation is supported by theoretical studies of motion extrapolation (ability to continue to predict accurately the position/speed of a moving object in the sudden absence of visual input) : Kaplan and colleagues implemented a realistic spiking recurrent neural network simulation of V1/MT (105), and studied cortical dynamics relaxation, when the coherent motion of a dot was abruptly interrupted by a “blank” period for three different lateral connectivity distributions (random, isotropic and anisotropic). The main finding, in agreement with this study, was that anisotropy was required to allow an efficient diffusion of motion information and achieve accurate motion/position extrapolation in the absence of FF validation (Fig. 4 in (105)).

#### 2.4. A neural correlate of perceptual bias in speed estimate for collinear motion

It is well established that “pop-out” and feature binding in Gestalt psychology do not depend invariably on attentive behavior, although they can be trained during perceptual learning and enhanced by top-down cognitive processes (11,15,16). The neural mechanisms, studied here in the anesthetized animal, can participate in automatic bottom-up attention, a form of sensory-driven selection that facilitates perception in the behaving state of a subset of the stimulus. Our working hypothesis is that the built-in bias in synaptic integration we report in primary visual cortex reflects lateral connectivity anisotropies engrammed during visuomotor development. This view is strongly supported at the functional level by the similarities in the visual cortical “association fields” found at the synaptic receptive field level in the anesthetized brain (our work), at the spiking discharge field level in the awake behaving state (Li and Gilbert) and at the behavioral and perceptual level (Field and Hess). We report here elementary changes at the subthreshold and spiking levels in V1, detectable at the intracellular level in the anesthetized state, that may not be strong enough to mediate an attention-guided behavior, but are efficient enough to produce an “automatic” pop-out effect.

Our electrophysiological findings constitute a potential correlate, as early as V1, of the human psychophysical speed-up bias in motion flow integration (47, 76). These previous psychophysical and computational studies from our lab showed that collinear AM sequences - composed of co-aligned ISO-Gabor patches along the global motion axis - are perceived by humans as moving “faster” than AM sequences composed by “parallel” CROSS-oriented elements, of the same exact physical speed. The speed bias decreased as the angle between the local GP inducer and the global motion axis increased. This bias was highly dependent on the orientation selectivity of the inducer and disappeared for blobs, which suggests a cortical origin. All reported features show a close link with the synaptic association field of V1 neurons (29) and the reported data.

Crucially, the speed for which the speed-up effect is the strongest in humans is 64 °/s. Assuming a parafoveal retino-cortical magnification factor ratio of 3 between human and cat (*<u>human</u>* : (106); *<u>cat</u>* : (107)), the optimal speed-up range of 40-96°/s seen in human would correspond to 120-288°/s in cat, which fits remarkably well the 100-312°/s reported here. The 64°/s optimal velocity for humans corresponds for the cat to a speed of 192°/s on the retina, and an apparent propagation speed of 0.19 mm/ms in cortex, in agreement with the inference made from our intracellular recordings. Most remarkably, the boosting mechanism revealed in this study confirms the prediction of an earlier computational model of our laboratory (47), which provided a simple conceptual framework where synaptic summation between feedforward and lateral AM inputs results in amplification and latency advance in V1 spiking responses.

Our intracellular study allows to go one step beyond this spike-based modeling. In (47), the composite synaptic summation is linear and the non-linearity of the functional effect is simply due to that of spike initiation. However, the intracellular study of the timing dependency of the association process suggests a non-linearity in the interaction process between feedforward (FF) and horizontal input, which becomes supra-linear when FF input lags by more than 5.5 ms the contextual signal (Fig. 7 C in (29); see also here Fig. 10, right pane)). It remains plausible that a minimal integration time is needed to recruit non-linear voltage-dependent mechanisms recruited by horizontal connectivity, such as the persistent sodium current or NMDA receptor activation (25, 108).

### 3. Concluding statements

In conclusion, the view presented here differs significantly from classical studies/models of top-down gain control of V1 processing which ignore the contribution *per se* of horizontal connectivity (11, 15, 16) or attribute the late enhancement of visual responses in V1 to the sole feedback from higher cortical areas (83). We do not argue here against the consensus that top-down signals are required, during attentive wakefulness, to express and/or amplify an integrative process intrinsic to V1. However, our recent demonstration of a “synaptic association field” in the anesthetized preparation demonstrates a preexisting structural bias intrinsic to V1, which could constitute the synaptic footprint of the neural architecture needed to implement Gestalt laws (29). This finding is further strengthened by a multiscale imaging study (40), suggesting that a critical threshold of spatial summation has to be passed in order to make these long-range interactions detectable, while temporal synergy conditions the effective binding of those spatial interactions (39). A likely interpretation is that some kind of resonance regime, such as produced here by the multiple stroke reafferent drive, is needed in the anesthetized state to propagate at the spiking level a functional bias already present in the subthreshold impact of horizontal connectivity.

The novel conceptual focus here is on the temporal features of horizontal propagation and on intra-V1 broadcasting. State-of-the-art modeling generally ignores the propagation distance-dependent delay component whose impact compromises greatly the analytical solving of mean-field equations (109); but see (110)). Interestingly, recent large-scale network simulations with topographically-organized horizontal monosynaptic connectivity and distance-dependent axonal conduction delays (0.10-0.60 m/s) demonstrate the consistency of spontaneous subthreshold waves with the asynchronous-irregular regime. The multiplexing of both processes defines a sparse-wave regime, more in line with a realistic view of a functional V1 (36). Although our interpretation of the present results seems in agreement with some of the computational literature on predictive coding applied to contour-through-motion extrapolation and trajectory extraction, they still remain to be confronted with alternative models which do not require propagation of a prediction. Other classes of probabilistic and static models (71) do not depend on propagation at finite speed and apply primarily to isotropic stimuli which are known to be suboptimal in activating V1.

Overall, our recordings suggest that fast collinear flow facilitates the refinement of anisotropy integration, distributed over the mesoscopic representation of trajectories across the “horizontal” layer plane of V1. The Surround-only CP-ISO anticipatory wave, captured “at the right position, at the right time” in a subset of our intracellular recordings, is compatible with a progressive topological refinement of activity diffusion in cortical space. In cat visual cortex, using annular stimuli and voltage sensitive dye imaging, Chavane et al reported correlates of “filling-in” responses (40), which fit with our subthreshold membrane potential recordings. Using intrinsic optical imaging (which might reflect more spiking than subthreshold activity), Chisum et al reported in the anesthetized tree shrew that ISO-oriented collinear Gabor arrays flashed simultaneously did enhance locally V1 sensory responsiveness (111). This latter study, which used a static input pattern of 2 second duration, activating simultaneously all the Surround sites, detected some activity strengthening in the areas directly (FF) activated by co-aligned local inducers. However, it failed to observe any spatial expansion of lateral activity in the cortical regions representing stimulus gaps. In contrast, our study predicts that the dynamic apparent motion animation of the same array, activating in succession the co-aligned local oriented inducers, would generate a continuous anisotropic propagation wave, most visible for AM speed faster than 50°/s. This diffusion process would, in turn, fill the gaps between local feedforward activation zones and promote the extraction of a motion trajectory riding on an expanding collinear contour. Evidence in V1 for an expansive “filling-in” process during apparent motion finds some support in fMRI imaging studies in humans (112). It could be tested spatio-temporally with more precision in cat and NHP, using voltage sensitive dye imaging and fast AM sequences.

Irrespective of the species-dependency of the visual spatial coverage of horizontal connections (7-8° in cats vs 1-2° in NHPs, in spite of similar axonal length), multiple stroke AM sequences appear best suited to dynamically recruit overlapping “lateral connectional fields” along the motion trajectory (90). The optimal separation in the visual field between the GP inducers might however be still species-specific, since the orientation selectivity of the horizontal facilitation process seems most prominent when co-stimulating the “Far” surround of GP-driven RFs.

This intriguing propagation of self-generated and propagated prediction at an early stage of visual cortical processing may underlie a more profound functional role. The classically reported “edge-detector” property of V1 neurons, as initially described by Hubel and Wiesel (46), highlights the fact that the strongest discharge is evoked by “parallel” elements swept towards the RF center orthogonally across its width axis. In contrast, at high AM speed (100-500°/s), the synaptic integration of the “silent” periphery was found to depend on the motion axis. This dynamic process unveils a spatial selectivity and anisotropy not present in the MDF mapped with static flashed light/dark impulses. This suggests that fast retinal flow produced by eye-movements could change radically contour integration selectivity of V1 neurons. The dynamic reconfiguration by fast retinal slip of the RF main axis could account for the peculiar geometry of saccadic scanpaths of human observers during the viewing of other human faces. Indeed, Alfred Lioukianovitch Yarbus long ago reported a striking similarity between the topographic layout of the saccadic scanpaths with the most salient long collinear contours outlining the perceptual skeleton of the face viewed by the observer ((27, 113, 114); review in (115)).

## Acknowledgments

We thank Vincent Bringuier, Frédéric Chavane, Jean Lorenceau, Peggy Seriès and Florian Gérard-Mercier for their past collaborative contribution to the concept of the “synaptic association field”. We are indebted to Frédéric Chavane, who participated in his PhD work to some of the early experiments. We thank Andrew Davison for helpful comments and help with the English. We thank Gérard Sadoc for the in-house development of visual stimulation and data acquisition software (Elphy^TM^), Kirsty Grant and Guillaume Hucher for histology and Aurélie Daret for her support during experimental surgery.

## Abbreviations

AM: Apparent motion
ASHP: Apparent Speed of Horizontal Propagation
CP: Centripetal
CF: Centrifugal
CROSS-axis: cross-oriented with the motion axis
CROSS-RF: cross-oriented (orthogonal) with the RF orientation preference
FF: Feedforward
GP: Gabor Patch
DN: dense noise
ISO-axis: collinear or parallel to the motion axis
ISO-RF: collinear or parallel to the RF orientation preference
MDF: Minimal Discharge Field
NHP: Non-Human Primate
RF: Receptive Field
RND: Randomized
SRF: Subthreshold synaptic Receptive Field
SN: sparse noise
V1: Area 17 in the cat, Primary Visual Cortex
WPSR: Wilcoxon paired signed rank statistical tests

## Author contributions

Y.F. conceived of the study.

Y.F., P.B., X.T., B.L.B., C.D., M.P., Y.P. and C.M. designed the experiments.

P.B. and M.P (“cardinal protocol”) and B.L.B., M.P., X.T. and C.D. (“radial protocol”) collected the intracellular data. P.B. (“cardinal” protocol) and B.L.B., C.D. and X.T. (“radial” protocol) implemented the data analysis. C.D. and M.P. designed the figures.

Y.F. and B.L.B. wrote the manuscript.

